# Gut-derived metabolites influence neurodevelopmental gene expression and Wnt signalling events in a germ-free zebrafish model

**DOI:** 10.1101/2021.07.06.451297

**Authors:** Victoria Rea, Ian Bell, Terence Van Raay

**Author notes:** Author contribution statement: VR and TVR conceived and planned the experiments; VR carried out WMISH experiments, RNA-seq analysis, qPCR. VR and TVR performed the lateral line screening; IB carried out KEGG pathway analysis and qPCR; VR wrote the manuscript with support from TVR; TVR edited the manuscript.

## Abstract

Small molecule metabolites produced by the microbiome are known to be neuroactive and are capable of directly impacting the brain and central nervous system, yet there is little data on the contribution of these metabolites to the earliest stages of neural development and neural gene expression. Here, we explore the impact of rearing zebrafish embryos in the absence of microbes on early neural development as well as investigate whether any potential changes can be rescued with treatment of metabolites derived from the zebrafish gut microbiota. Overall, we did not observe any gross morphological changes between treatments but did observe a significant decrease in neural gene expression in embryos raised germ-free, which was rescued with the addition of zebrafish metabolites. Specifically, we identified 361 genes significantly down regulated in GF embryos compared to conventionally raised embryos via RNA-Seq analysis. Of these, 42 were rescued with the treatment of zebrafish gut-derived metabolites to GF embryos. Gene ontology analysis revealed that these genes are involved in prominent neurodevelopmental pathways including transcriptional regulation and Wnt signalling. Consistent with the ontology analysis, we found alterations in the development of Wnt dependent events which is rescued in the GF embryos treated with metabolites.

## INTRODUCTION

Animals and microbes share a deep evolutionary history as animal development emerged and co-evolved with a microbe-rich environment^1^. The gut microbiome codes for biochemical functions that host genomes cannot encode, such as the breakdown of otherwise indigestible macromolecules into products that their hosts can utilize^2^. The microbiome has been implicated in neural development and function, and consequently, perturbation of the microbiota is implicated in neurological disease^3–6^. It is known that metabolites act as the communication signals between host and microbiome in the form of neuromodulators or neurotransmitters^7^. Both neural and circulatory routes have been proposed as a means of gut-brain signaling including the vagus nerve and enteric nervous system (ENS), and direct absorption from the intestinal lumen into the blood stream^8,9^. The vagus nerve and ENS are sensitive to gamma amino butyric acid (GABA), serotonin, histamine, and acetylcholine, all of which are produced by the gut microbiota^9^. Small molecules such as short chain fatty acids (SCFAs) produced by the gut microbiota, can enter the blood stream via the intestinal lumen, and cross the blood brain barrier (BBB) where they can then interact with the brain and affect neural transmission^10^. Therefore, the correlation between the gut microbiome and the brain is unlikely due solely to the presence of bacteria, but more likely due to the metabolites and small molecules that bacteria produce as fermentation products. Studies have shown that metabolites are critical signalling molecules produced by bacteria and utilized by the host, yet there is limited data on the contribution of gut-derived bacterial metabolites on the earliest stages of neurodevelopment. Here, we use zebrafish neurodevelopment as a proxy for evaluating the contribution of gut-derived microbe metabolites to early neural development and gene expression.

## RESULTS

### Microbes are necessary for timely neural gene expression

To determine if microbes are required for neural gene expression and patterning, the spatial distribution of select neural genes were analyzed using whole mount in situ hybridization (WMISH) in conventionally raised (CV) and germ-free (GF) zebrafish embryos. All embryos in each cohort were raised in parallel, were time and stage matched, randomly assigned in the WMISH protocol, and processed in parallel to ensure that differences in gene expression were not due to an offset in overall development or procedure. The WMISH data demonstrated a significant decrease in expression of five out of six target genes in germ-free embryos at 2 days post-fertilization (dpf) (Fig. 1A). All target genes, with the exception of *isl1*, showed a decrease in relative level of expression. However, expression of *notch1b, ngn1* and *ascl1a*, which had reduced expression levels at 2 dpf, were increased in 4 dpf germ-free embryos, suggesting a delay in expression of these genes under germ-free conditions (Fig. 1B). Expression levels of *fgf8* and *phox2bb* remain decreased in the GF group at 4 dpf relative to CV controls while *isl1*, which showed little difference between treatment groups at 2 dpf, showed a significant decrease in expression in the GF group at 4 dpf. Interestingly, it is the genes that are more ubiquitously expressed that display a delay in expression rather than an overall decrease, yet there are no gross morphological differences between conventionally raised and germ-free zebrafish (Fig.1F). Taken together, this suggests that there is a delay in neural development in the absence of microbes.

**Figure 1.**
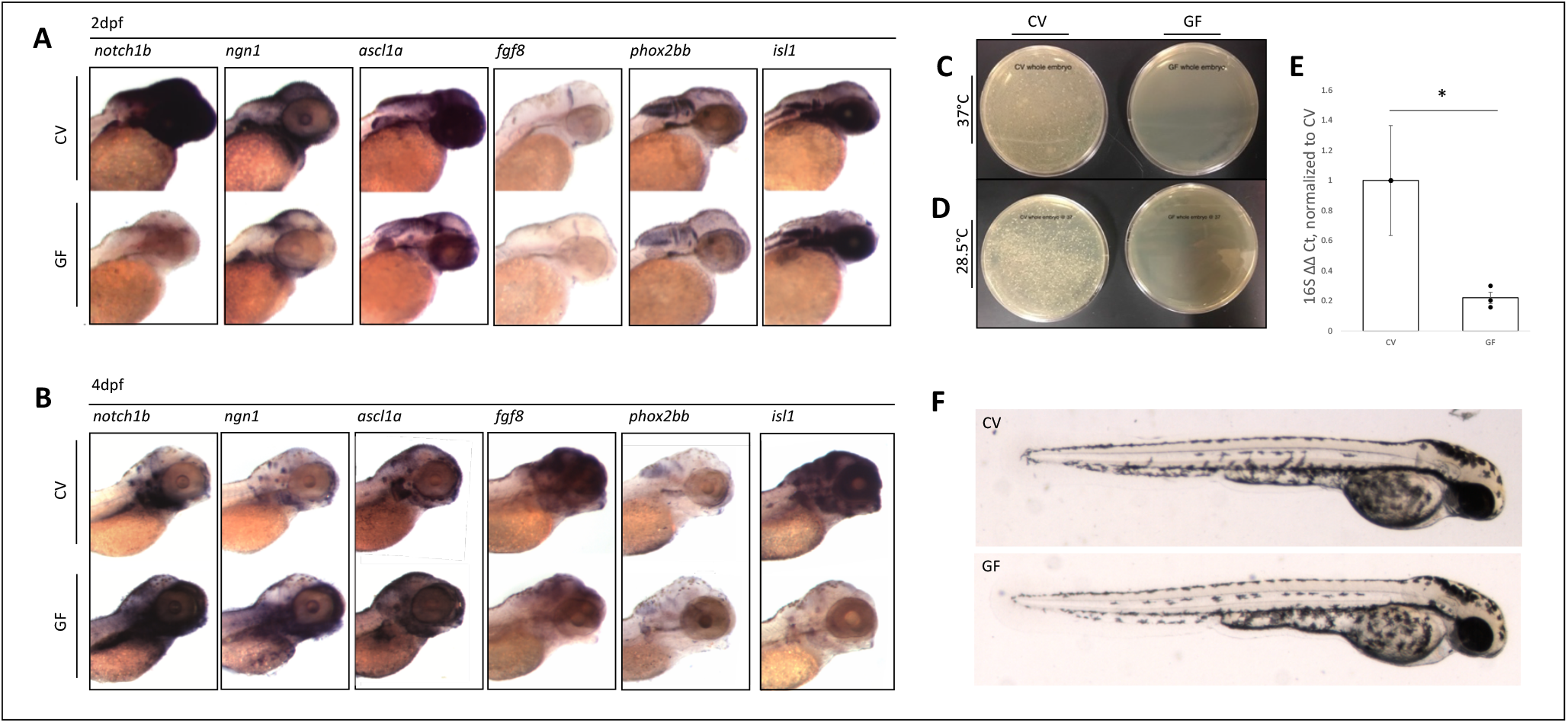
Microbes are necessary for timely neural gene expression. **A)** WMISH of target genes in conventionally raised (CV) and germ-free (GF) embryos at 2dpf. RNA expression of target genes *notch1b*, (N = 6, 108 embryos) *ngn1* (N = 3, 54 embryos), *ascl1a* (N = 2, 36 embryos), *fgf8* (N = 6, 108 embryos) and *phox2bb* (N = 1, 18 embryos) is reduced in the absence of microbes at 2dpf. Expression of *isl1* (N = 3, 54 embryos) shows little difference between groups. **B)** WMISH of target genes in conventionally raised and germ-free embryos at 4dpf. RNA expression of target genes *notch1b* (N = 4, 72 embryos), *ngn1* (N = 2, 36 embryos), and *ascl1a* (N = 1, 18 embryos), show an increase in expression in the GF group compared to their CV counterparts at 4dpf. Expression of *fgf8* (N = 2, 36 embryos) and *phox2bb* (N = 2, 36 embryos) remains reduced in comparison to the CV group. **C-D)** Whole homogenized single CV (left) or GF (right) embryos plated on brain heart infusion media and left at **C)** 37°C or **D)** 28.5°C for 24 hours. **E)** RT-qPCR analysis of universal 16S rRNA gene in CV and GF embryos (* = p < 0.05 in a one-way ANOVA, based on delta, delta Ct), normalized to ef1*α*. **F)** Live images of 2dpf zebrafish embryos for morphological comparison. N values represent biological replicates.

### Lack of microbes results in global decrease in neural gene expression

The general decrease in the majority of our WMISH probes suggests that microbes and/or their metabolites might have a more general role in neural development. To determine this, we performed RNA-Seq analysis on RNA enriched from zebrafish heads under three different conditions. As above, we analyzed the gene expression in zebrafish embryos that were conventionally raised and germ free. To determine if bacterial metabolites were sufficient to affect gene expression, we treated GF embryos at shield stage to 60% epiboly with metabolites isolated and filter sterilized from adult zebrafish guts (ZM). Total RNA was extracted from the heads of zebrafish embryos at 2 dpf, the height of neurogenesis, enriched for mRNA and subjected to RNA-seq analysis.

Our first observation was that differential gene expression analysis revealed a general decrease in gene expression in the GF group (Fig. 2A) with over 2000 genes displaying some decrease in expression in the GF group compared to CV. Secondly, we observed a substantial decrease in the variation of expression in GF compared to the other treatments (Fig. 2B). Importantly, ZM treatment sufficiently rescued gene expression in GF larvae, along with an increase in the variability (Fig. 2B,C). Taken together, this suggests that in the absence of metabolites, gene expression is uniformly maintained at a seemingly basal level and that metabolites are both necessary and sufficient to elevate or enhance gene expression.

**Figure 2.**
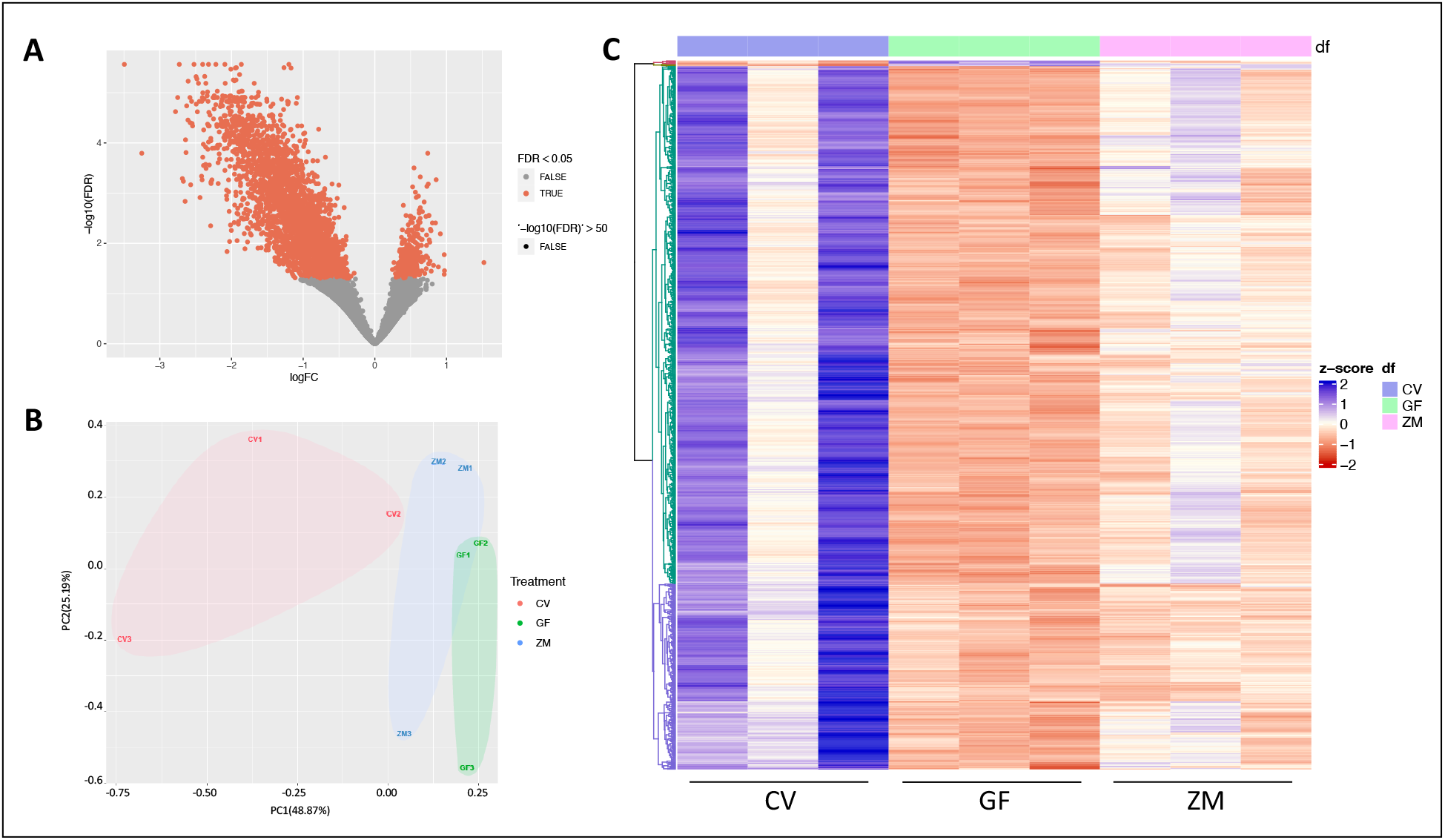
Microbes both necessary and sufficient for general gene expression in the developing nervous system. **A)** Volcano plot comparing DEGs between germ-free larvae and conventionally raised larvae. **B)** Principal component (PC) analysis of count data from CV, GF and ZM zebrafish larvae at 2dpf. **C)** Heatmap of top 1000 differentially expressed genes between 3 biological replicates of CV, GF and GF treated with zebrafish gut metabolites (ZM) embryos at 2dpf. Generated with DeSeq2 and ComplexHeatmap.

In order to look more specifically at the biological processes and molecular functions associated with germ-free treatment, gene ontology (GO) analysis was performed on the subset of genes whose expression was down regulated at least two-fold. Absence of microbiota resulted in a significant decrease in expression of 354 genes (log fold change < -2, FDR <0.05) (Supplemental Table 1). GO statistical overrepresentation tests revealed that these genes are largely involved in RNA binding, DNA binding and modification, transcription regulation, neurogenesis, axonogenesis, and Wnt signaling (Supplemental Table 2). It should be noted that there were also seven genes upregulated in the GF group compared to CV, however these genes did not have any biological significance in statistical overrepresentation tests.

### Metabolites are sufficient to rescue the expression of neural development genes

The addition of metabolites to germ-free zebrafish rescued the expression of numerous genes that were significantly down regulated in GF (p <0.05, FDR <0.05) compared to CV larvae, although not to the extent observed in CV larvae. We considered gene expression to be rescued by zebrafish metabolites if the log fold change of a gene in ZM-GF was in the opposite direction of the log fold change of the same gene in GF-CV (GF-CV FDR <0.05, ZM-GF FDR <0.1). Using these criteria, the expression levels of 42 genes were rescued by metabolites (Fig. 3A). That is, 39 genes were down regulated in the GF group compared to CV but up regulated in the ZM group compared to GF, and 3 genes were upregulated in the GF group compared to CV and downregulated in the ZM group compared to GF (Fig. 3A,C). The expression levels of these 39 upregulated genes were highly variable between the 3 CV biological samples (Fig. 3H), consistent with the analysis of the entire data set (Fig. 2B, C). Interestingly, this variation was considerably reduced in the GF samples, but increased again upon treatment with metabolites. We analyzed the function of these 39 genes using DAVID, an online bioinformatics tool that condenses gene lists and associated biological terms for functional annotation using four analysis modules: Annotation Tool, GoCharts, KeggCharts, and DomainCharts (https://david.ncifcrf.gov/). The output from DAVID was plotted via ComplexHeatmap v2.5.5 package for R (Fig. 3B, Supplemental Table 3). Of the 39 downregulated rescued genes, 30 had biological significance in a gene function analysis in DAVID (Fig. 3B). The genes rescued by metabolites are largely involved in cellular processes related to DNA binding, nuclear import, transcriptional regulation, and mRNA splicing; as well as neural developmental processes involving Wnt signalling and axonogensis. Overall, this emphasizes the importance of metabolites during early neural development. The three genes that were down regulated and rescued; *cryba2a, crygmxl2* and *crybm2d20*, are associated with eye and eye lens development and were not included in the DAVID plot but are included in the normalized count plot (Fig. 3C). Curiously, these 3 genes had significantly higher levels of expression in CV compared to the others and the changes in expression, while significant, were to a smaller degree compared to the genes whose expression were upregulated in ZM. To validate the RNA-seq data, select genes from this list were quantified via RT-qPCR from independent sources of mRNA for the three conditions. (Fig. 3C-G). These results support both the RNA-seq and WMISH data that metabolites are both necessary and sufficient for gene expression.

**Figure 3.**
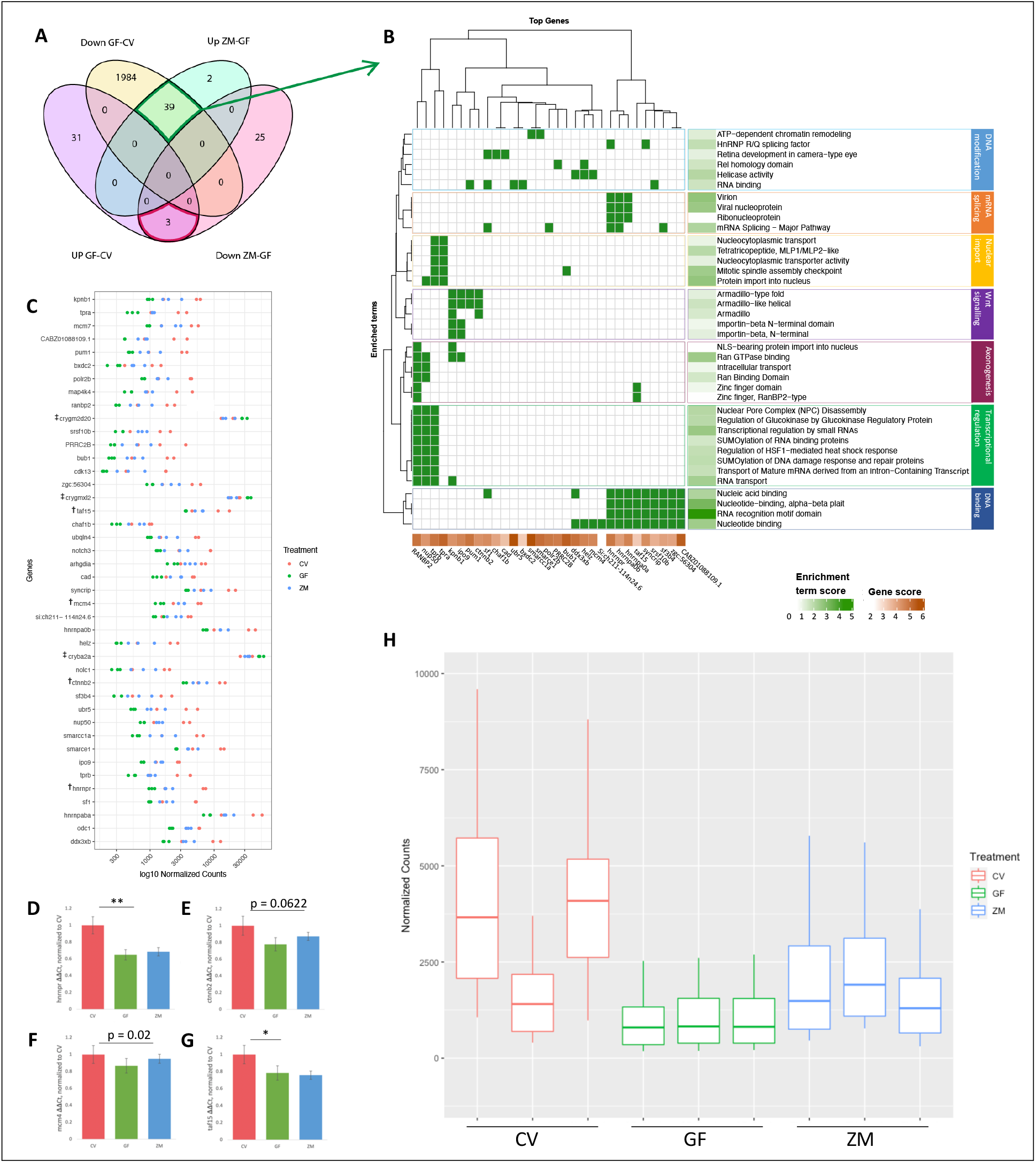
Metabolites are sufficient to rescue neural gene expression in GF larva. **A)** Venn-diagram comparing gene expression levels between CV and GF and those rescued by the addition of zebrafish metabolites to GF embryos (ZM-GF) (p < 0.05, GF-CV FDR < 0.05, ZM-GF < 0.1) **B)** DAVID generated plot of 31 of the 39 rescued genes and their associated enrichment terms (note: 8 genes did not contribute to significant over representation in DAVID output). Enrichment terms largely fall within seven major biological processes. **C-G)** Count comparison and RT-qPCR validation **(D-G)** of four rescued genes (marked with^**†**^) from the RNA-seq dataset. **C)** Normalized counts of all 42 genes (39 downregulated plus 3 upregulated, noted by ^‡^) whose expression was rescued with metabolite treatment. **D-G)** RT-qPCR validation of *hnrnpr* **(D)**, *ctnnb2* **(E)**, *mcm4* **(F)** and *taf15* **(G)** (* = p < 0.05, ** = p < 0.01; One way ANOVA of all 3 groups standard weighted means analysis, 3 independent samples, 2 degrees of freedom, total p value is as stated). **H)** Boxplot of normalized counts of 39 downregulated and rescued genes between treatment groups with outliers removed. Top and bottom of box represents the 75^th^ and 25^th^ percentile respectively. The 50^th^ percentile and solid horizontal line in the box represents the median. Whiskers represent largest and smallest value within 1.5 times interquartile range above 75^th^ percentile and below 25^th^ percentile respectively.

### Neural development is disrupted in germ-free embryos

The absence of a gross morphological phenotype but significant decrease in gene expression in GF prompted us to investigate the consequence of being germ free at the cellular level. Using acetylated *α*-tubulin immunostaining as a general axon marker in combination with transgenic Glial fibrillary acidic protein (GFAP:GFP) to mark neural stem cells and glia, we looked at the general architecture of the zebrafish larval brain at 2 dpf. Consistent with our WMISH and RNA-seq data, we observed a modest and generalized disorganization of neurons and glia in GF larvae. In particular, we observed an uneven distribution in GFAP-GFP fluorescence in GF compared to CV and ZM treatments (Fig. 4 D-F). Upon closer inspection of rhombomeres in the hindbrain, GFAP:GFP fluorescence reveals changes in the pattern of rhombomeres in GF embryos compared with CV and ZM treatments (Fig. 4J-L).

**Figure 4.**
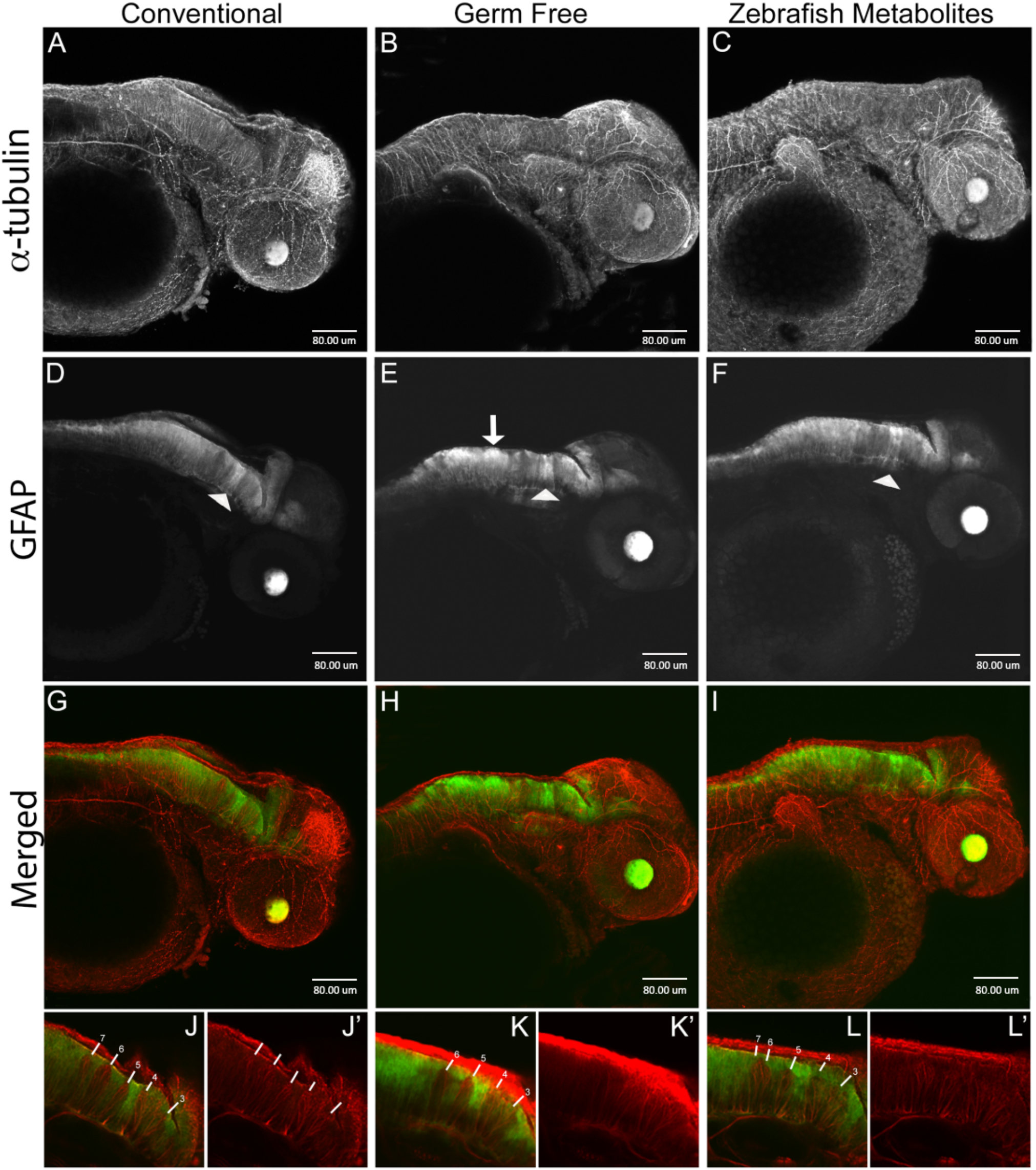
Neural development is disrupted in germ free embryos. Confocal projection images of zebrafish embryos at 2dpf. (A-C) *α*-tubulin immunostaining, (D-E) GFAP fluorescence. GFAP:GFP fluorescence (**D-F**) displays a non-uniform distribution in the hindbrain in germ free embryos (white arrow in E) and to some extent in ZM treated embryos. The white arrowheads identify the GFAP tract between rhombomeres 4 and 5 which does not appear to be significantly altered in germ free embryos. **G-I**) Merged images of *α*-tubulin and GFAP:GFP. **J-L)** Representative single layer images of regions in the hindbrain. In conventional embryos, rhombomere tracts, 3-7 are readily identifiable by the relative absence of GFAP fluorescence. The higher intensity staining between rhombomeres 4 and 5 provides a landmark for their easy identification. Note the absence of rhombomere 7 in germ free embryos, and the seemingly merged tract 6 and 7 in ZM treated embryos. More examples are presented in supplementary figures 5 and 6.

The zebrafish lateral line is a mechanosensory organ that requires coordination of cell proliferation, migration, and differentiation^11^. Furthermore, it has been well documented that Wnt signalling plays an active role in primordial neuromast deposits of the lateral line^11^. The posterior lateral line of the trunk arises from the first placode at around 18 hpf, which gives rise to neuroblast precursors and the first primordium that migrates down the trunk over the next 20 hours depositing cellular rosettes that eventually differentiate into neuromasts^12^. Thus, investigating the lateral line in GF larva would be a useful way to evaluate neural cell migration and specification with potential links to Wnt signaling, which we observed to be perturbed in our RNA-seq analysis in GF larvae. To investigate this, we first looked at neuromasts by WMISH with *ascl1a, notch1b* and *isl1* (Fig. 4A-C; Supplemental Figure 1) at 4 and 5 dpf. We observed alterations in the location and number of primordial neuromasts of the lateral line in GF larvae, which was partially rescued with ZM (Fig. 4A-C). To further Investigate the neuromasts of the lateral line, we performed live vital dye analysis with Diasp and DiOC6, which further demonstrated that development of the posterior lateral line is disrupted in germ-free embryos and rescued to some extent in the embryos treated with zebrafish metabolites (Fig. 4D-I, Supplemental Fig. 2,3). At 3 dpf, neuromasts of the posterior lateral line in the trunk of GF embryos appear unevenly distributed, more anteriorly positioned and immature compared to the CV and ZM embryos, where on average, more of the neuromasts in both the CV and ZM groups have migrated past the anal pore, consistent with the WMISH data. Further, the terminal neuromasts are less well-developed and in some cases missing in GF embryos (Fig. 5G-I, Supplemental Fig. 3). We also observed changes in posterior lateral line neuromasts at 4dpf, where GF embryos had between one and four trunk neuromasts compared to CV embryos that had between four and seven (Supplemental Fig. 4)

**Figure 5.**
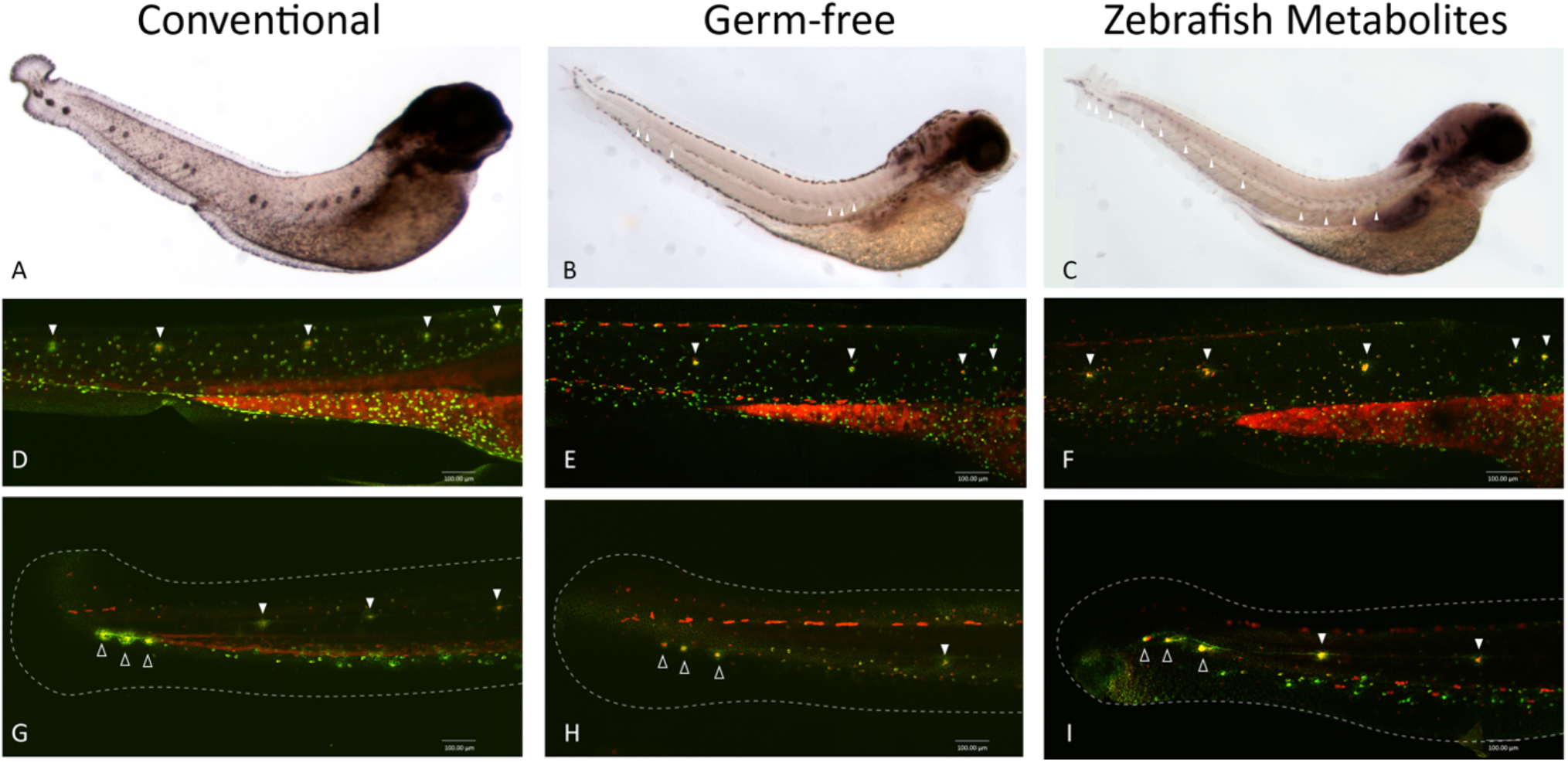
Posterior lateral line development is disrupted in germ-free embryos. **A-C)** WMISH of *isl1* in 4dpf embryos. Larvae from each treatment group were processed in parallel. CV larvae were overstained to allow comparison to GF and ZM groups which have reduced staining, but visible neuromasts (white arrowheads). **D-F)** Trunk neuromasts of the posterior lateral line in 3dpf embryos incubated in a mixture of vital dyes Diasp and DiOC6 to identify hair cells (red) and accessory cells (green) of the lateral line. Neuromasts are marked with white arrowheads. The intense staining in the yolk extension provides a useful position reference. **G-I)** Posterior trunk neuromasts (solid white arrowheads) and terminal neuromasts (outlined arrowheads) in the tail identified with Diasp and DiOC6 in 3dpf embryos. Tail fins are outlined with dashed line for reference. Terminal neuromasts appear to be smaller and less well-developed in GF embryos. Some neuromasts of the posterior tail, as well as some terminal neuromasts, are missing in GF embryos. More examples are presented in supplementary figures 2-4.

### Metabolites affect Wnt signalling

The combination of the DAVID output (Fig. 3B, 6A) identifying Wnt signaling and the effect of neuromast development (Fig. 5), a Wnt dependent event, as being affected by bacterial metabolites prompted us to further investigate Wnt signaling. Wnt signaling is important in many developmental processes including cell fate determination, proliferation, and axonogenesis and migration^13^. Indeed, there is evidence that bacteria activate Wnt signalling to regulate the inflammatory response^14^. Further, several studies have demonstrated that bacteria activate Wnt signalling with effects on the intestinal epithelium^15–18^, reproductive tract^19,20^, and respiratory tract^21,22^. Studies in both mice and zebrafish have shown that bacteria induce intestinal cell proliferation in a Wnt dependent manner and that germ-free animals have decreased Wnt signalling and decreased intestinal epithelial cell proliferation^23,24^. To explore this further, we identified 75 genes that the Wnt community has identified as being targets of, or important in, Wnt/β-catenin signaling (The Wnt Homepage; Fig. 6B). We found that 25 of the 75 genes exhibited reduced expression in GF and rescued expression in ZM pattern. We validated two of these genes (*sp5a* and *ctnnb2*, Fig. 6C, D) and further performed a KEGG analysis (Fig. 6E), all of which demonstrates that the Wnt pathway is one of the major signaling pathways affected by bacterial metabolites. In addition to Wnt signaling, other developmental signaling pathways were also affected, including TGFβ, Hedgehog and Notch (Supplemental Fig. 7), consistent with the broad decrease in gene expression in the GF treatment.

**Figure 6.**
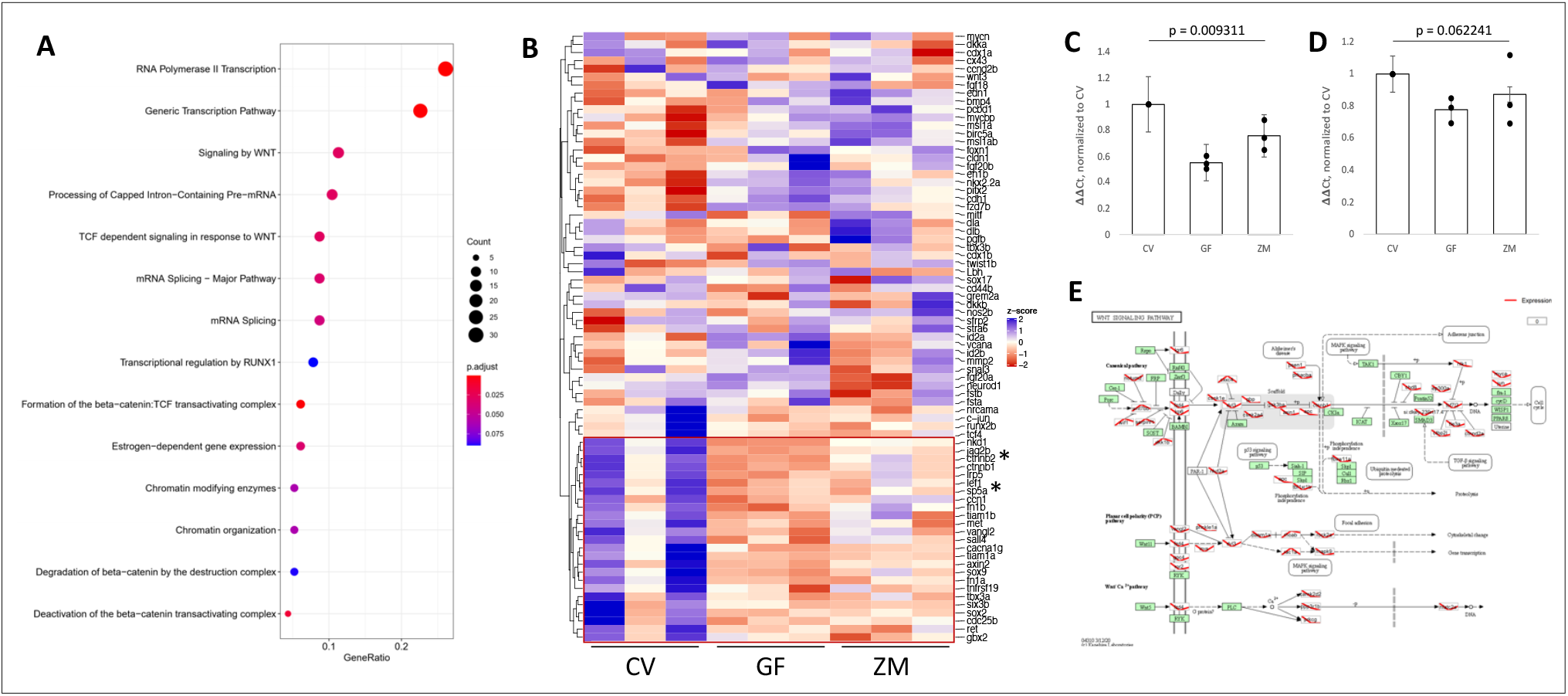
Metabolites Affect Wnt signaling. **A)** Enriched pathway analysis of the downregulated genes in the GF group via KEGGprofile. **B)** Heatmap of target genes of Wnt/β-catenin signaling identified by the Wnt signalling community. Box in red shows genes that are affected in our dataset. Asterisks identify two genes that were validated by RT-qPCR (**C**, *sp5* and **D**, *ctnnb2;* One way ANOVA of all 3 groups standard weighted means analysis, 3 independent samples, 2 degrees of freedom, total p value is as stated). **E)** KEGG profile output of Wnt pathway and genes from complete RNA-Seq dataset. Expression levels are shown in red where the left side (CV) is arbitrarily set to 0, the middle point is GF and the right point is ZM.

Because Wnt signalling was affected in our dataset, we investigated whether the decrease in expression of developmental genes was at least in part due to down regulated Wnt signalling. We used two compounds known to affect Wnt signalling to treat CV and GF embryos and analyzed expression of neurodevelopment gene *ascl1a* and Wnt target *axin2* via WMISH. Conventionally raised embryos were treated with XAV939, a small molecule that inhibits Wnt activity by inhibiting tankyrase (TNKS) and β-catenin mediated transcription^25,26^. GF embryos were treated with BIO, a compound that functions as a Wnt activator through inhibition of GSK3β^27^. Each compound was added to either CV or GF embryos, respectively, immediately after the GF embryos were sterilized and all four groups of embryos were allowed to develop to 2 dpf and processed in parallel. Both the GF and the CV + XAV939 treated larvae displayed a relative decrease in expression of both *ascl1a* and *axin2*, consistent with our previous findings. Importantly, the GF + BIO treated larvae displayed relatively higher expression like that of the CV larvae (Fig. 7). Spatially, expression was predominantly affected in the hindbrain (Fig. 7A-D) and the posterior recess of the hypothalamus^28,29^ (Fig. 7E-H). Indeed, specifically inhibiting Wnt appears to have the same effect on expression of *ascl1a* and *axin2* as deriving the embryos germ-free. Further, treating GF embryos with a Wnt activator rescues the expression of these genes to a level that is comparable to CV larvae. Taken together these results suggest that Wnt signalling is dependent on microbes at some level, though more research is necessary to determine causation.

**Figure 7.**
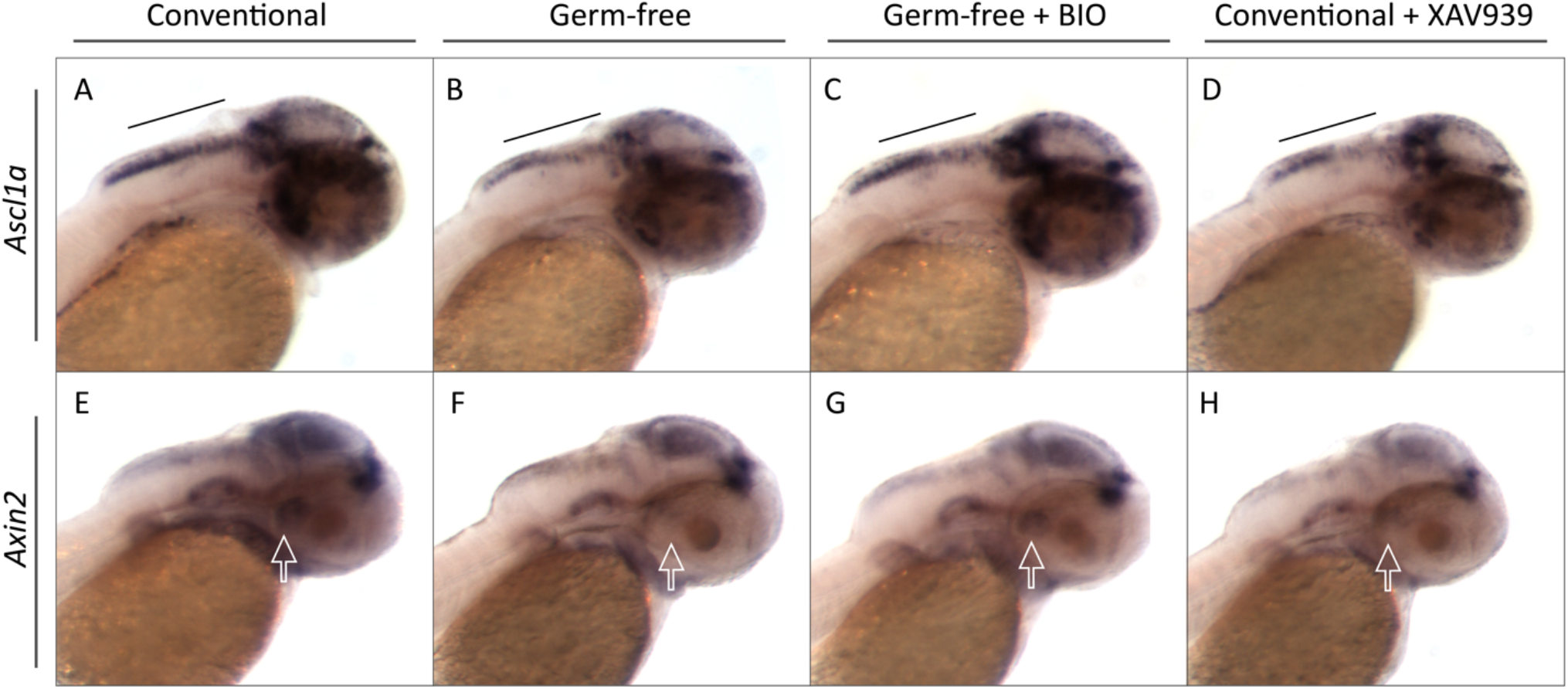
Specific regulation of Wnt signaling mimics the GF and ZM conditions. Representative images of whole mount *in situ* hybridization on 2 dpf conventional embryos, germ-free embryos, germ-free embryos treated with known Wnt activator BIO, and conventional embryos treated with known Wnt inhibitor XAV939 for genes *ascl1a* (A-D) and *axin2* (E-H). Black bar identifies the hindbrain region where there is less staining in both the GF and conventional treated with Wnt inhibitor than the CV or GF treated with Wnt activator. Solid white arrows represent the posterior recess of the hypothalamus^28,29^.

## DISCUSSION

In this study, we evaluated the contribution of bacteria and gut-derived metabolites on neural gene expression and development. Making zebrafish germ free from 0 to 4 days appeared to have no gross morphological effect at early larval stages and these fish developed to adulthood identical to their conventional and metabolite treated cohorts (data not shown). In other models, germ-free animals initially appear normal but tend to function at a lower metabolic efficiency ^30–32^ and have negatively impacted development of other organs and organ systems^4,33–35^. Intestinal microbes provide significant biochemical functions to generate metabolites that eukaryotes are incapable of generating such as butyrate, propionate, and acetate^2^. While there may be no gross morphological effects, we do demonstrate that gut-derived metabolites are in large part responsible for regulating critical signalling pathways in the brain, especially during neural development.

### Large Genomic Effects

Overall, we observed a general decrease in expression of many genes in GF, which was partially rescued by zebrafish metabolites. Further, we found significant expression level variability in the CV and ZM groups, which was dramatically reduced by making larvae germ-free. This suggests that there is a basal level of expression that is amplified by bacterially derived metabolites. It is interesting that we did not observe gross morphological differences between the treatment groups, which we attribute to the maternal contribution of metabolites in the yolk. These maternally-derived metabolites may also contribute to the basal level of gene transcription that we observed. Nonetheless, given that GF gene expression can be rescued by the addition of metabolites provides an attractive platform in which to study the contribution of purified or specific metabolites to biological processes as has been observed in GF mouse models^3^. Further, the hair cells in the lateral line are analogous to mammalian inner ear hair cells and as such provides a tractable model for understanding the contribution of bacterial metabolites to relevant biological processes^36^. We are currently identifying other biological processes that are perturbed in GF larvae and rescued by metabolites to further pursue this model.

### Mining the contributions of metabolites

Rearing zebrafish embryos germ-free resulted in a significant reduction in the expression of 354 genes, and an increase in seven genes. Treating GF embryos with metabolites derived from the zebrafish gut significantly rescued the expression of 42 of these genes. Using DAVID analysis, we found that RNA binding, DNA binding and modification and transcription regulation genes were the major genes being affected by both the absence and addition of metabolites. While the levels of transcription in ZM larvae did not reach CV levels, they were sufficient to rescue defects in the developing nervous system caused by being germ-free. Our findings are consistent with other studies that have demonstrated that microbiome depletion is linked to alterations in RNA processing, particularly alternative splicing^6^, and previous studies demonstrating that gut microbiome metabolites can affect DNA and RNA binding, processing, and transport^37–39^.

### Wnt Signaling/Lateral line

We observed that several prominent developmental signaling pathways are responsive to gut metabolites, most notably Wnt signaling. Indeed, when we enriched for Wnt signaling genes identified by the Wnt community, we found a significant reduction in 25% of these in GF larvae. Wnt signaling is well-recognized for its role in neural development^13^, posterior lateral line^11,40^ and mental disorders^41^. Interestingly, we found similar results via WMISH of *axin2* in 2 dpf GF embryos and CV embryos treated with a Wnt inhibitor as a recent report of hypothalamic genes associated with Wnt signaling and anxiety in a zebrafish Lef1 mutant^28^. As Wnt signaling is also influenced by bacteria ^14^, it is not surprising that we observed alterations in Wnt signalling dependent processes. Wnt dependent activities, such as the migration and development of the lateral sensory hair cells, were affected in GF and rescued in ZM. The uniform distribution of GFAP in CV larvae was also disrupted in the GF treatment and rescued by the ZM treatment. GFAP is a marker of neural stem cells and glia and we observed an increase in GFAP:GFP fluorescence in GF larvae, which is consistent with the delay in neurogenesis that we observed by WMISH and seen in Wnt1 morpholino knockdown studies^42^

Independent studies have also demonstrated that Wnt signaling was downregulated in germ free mice, which displayed defects in thalamocortical axonogenesis and aversive somatosensory behaviours^3^. Further, the Wnt/β-catenin effector Lef1 is required for the development of the hypothalamus and differentiation of anxiolytic hypothalamic neurons in both zebrafish and mice, which also displayed increased anxiety in zebrafish in the absence of Wnt/β-catenin signaling^28^. A preliminary touch response assay revealed that GF larvae swim farther and more quickly after stimulus with a probe than CV larvae (Supplemental Fig. 8), which is in line with the Lef1 zebrafish mutant and other reports of locomotor activity in GF zebrafish^28,43,44^. Taken together, there is strong evidence that metabolites are directly regulating Wnt signaling, which impinges on several neurodevelopmental pathways

### Comparison to other studies

Our expression results are consistent with previous reports in microbiome depleted mice. A recent study by Vuong et al. (2020), found that microbiome depletion altered the expression of 333 genes in the brains of embryonic mice, including many genes involved in axonogenesis. We found 67 of the same genes differentially expressed in GF zebrafish embryos. Somewhat surprisingly, one of the genes rescued by metabolites in both the Vuong et al., (2020) study and in the current analysis is *ctnnb2*, CTNNB1, the central contributor to the Wnt signalling pathway, which has been implicated in other studies looking at specific microbial species^16,45,46^. Independent of germ-free status, both Wnt signalling and axonogenesis have been implicated in studies of the microbiome^3,14,24,47^. We also found a substantial overlap between differentially expressed genes in the current dataset and genes identified as candidate risk genes for neurodevelopmental disorders, where 256 genes that were downregulated in GF larvae compared to CV larvae are orthologous to genes identified by SFARI (Supplemental Table 4). The independent and consistent identification of Wnt signaling as a target of bacterial metabolites, the well-established role of this pathway in neural development, and the role this pathway plays in so many diseases, elevates this pathway to a new level. Further, comparison of germ-free animal models should ultimately identify a universal set of genes most likely affected by metabolites.

### Conclusions

It is becoming quite clear that neural development does not occur in a sterile and metabolite-free environment. However, understanding how these metabolites impinge on neural development is still in its infancy. Consistent with other independent investigations, we identified significant changes in neural gene expression that are under the influence of bacterially-derived metabolites. With such substantive changes, it can be difficult to identify the most important players, but the Wnt signaling pathway has emerged as playing a leading role in this process. Given that this pathway first arose in multicellular eukaryotes and plays such a significant role in development and disease, perhaps it shouldn’t be surprising that its regulation co-evolved with the bacterial colonization of multicellular eukaryotes. Further investigation into the metabolite-Wnt-neurodevelopment axis could ultimately lead to better therapies for the myriad of Wnt-related mental disorders^41^.

## METHODS

### Zebrafish maintenance

Zebrafish from the standard wild-type Tübigen (TU) line were raised and maintained in accordance with the Animal Protocol Utilization # 3614 using standard protocols^48^. Zebrafish were maintained on 14:10 hour light: dark cycle. Larvae were obtained by natural spawning and cultured in zebrafish embryo medium (EM; 0.00006 w/v% Instant Ocean® Sea Salt solution and 0.0001% methylene blue in purified distilled water) at 28.5°C. For *in vivo* imaging and head dissection, larvae were anesthetized with 0.04% tricaine.

### Generation and treatment of germ-free larvae

Larvae are collected within two hours of fertilization and develop in a 28.5°C incubator. At shield stage to 60% epiboly (specification of the 3 germ layers but before neurogenesis) larvae are divided into CV and GF groups. CV larvae are left at room temperature (RT) while the GF group is sterilized at RT to normalize their development. GF larvae are immersed in filter sterilized Gentamicin (100*μ*g/mL) for one hour and subsequently washed in 0.003% hypochlorite followed by three five-minute washes in sterile embryo medium. Embryo treatment is performed under a laminar flow hood to ensure sterility. Post sterilization, larvae from both groups are placed in a 28.5°C incubator. After 24 hours, a 20µL sample of both EM and a single homogenized embryo are plated on separate brain heart infusion (BHI) agar plates, a non-selective, nutrient rich growth medium, along with an empty control plate (exposed concurrently with samples) and incubated at 28.5°C or 37°C for 24 hours to test for sterility. Upon confirmation of sterility (0 visible colonies), larvae are harvested at the appropriate time points outlined below.

### Whole mount *in situ* hybridization (WMISH)

Zebrafish larvae for WMISH were treated with sterile PTU (0.003%) at 24 hpf to reduce pigment development, harvested at 2 or 4 dpf, manually dechorionated and immediately fixed overnight in 4% paraformaldehyde (PFA, Sigma) in 1X PBS. WMISH was performed as previously described^49^. DIG-labelled probes were synthesized by *in vitro* transcription (New England BioLabs Inc.) with appropriate polymerases, following the manufacturer’s instructions and after plasmid linearization with appropriate restriction enzymes.

### Imaging

WMISH-stained larvae were mounted in 100% glycerol. Live larvae were anesthetized in 0.04% tricaine, embedded in 2% methyl cellulose and imaged with dissecting microscope (V8 Zeiss) mounted with a X camera and imaged using Q-Capture software (v 3.1.3.10). Fluorescent images were captured using a Leica CLSM SP5 confocal microscope using LAS AF imaging software v2.7.7.

### Zebrafish metabolites

#### Extraction

Pools of ten adult male zebrafish were euthanized in an ice bath slurry for at least 10 minutes according to standard procedures^50^, followed by surgical removal of the intestine. Intestines were resuspended in sterile 1X PBS at a 1:3 weight:volume ratio (∼1 mL) and vortexed for approximately 1 minute to resuspend intestinal contents followed by centrifugation at 14,000xg for 30minutes. The supernatant was filter sterilized through a 0.22*μ*M filter and stored at -20°C.

#### Treatment

Germ-free larvae were immediately treated with undiluted zebrafish metabolites added directly into the sterile embryo medium by adding 200uL (equivalent of 2.7 adult guts worth) of metabolites mixed with 15mL EM in a 10cm sterile dish containing ∼100 larvae at ∼60% epiboly. After 24 hours, a 20µL sample of both EM and a single homogenized embryo were tested for sterility as described above.

### RNA sequencing and analysis

At 2 dpf, larvae where euthanized in 0.04% Tricane and the heads were surgically removed from the body at the base of the hindbrain. RNA was extracted from a pool of five heads for each treatment using the GENEzol™ TriRNA Pure Kit (Froggabio). RNA samples were DNase treated using the Invitrogen™ DNA-free™ DNA Removal Kit (Thermo Fisher Scientific). An RNA integrity number (RIN) of more than 8.0 was confirmed for all samples using the 4200 Tapestation system (Agilent). Poly(A) mRNA was prepared using the NEBNext® Ultra™ II Directional RNA Library Prep Kit for Illumina® (New England BioLabs) and 2 × 100bp paired-end sequencing at a depth of 80-100 million reads per sample was performed using the Illumina Novaseq 6000 platform by the University of Toronto Donelly Sequencing Centre. FastQC v0.11.8 and HISAT2-2.1.0^51,52^ were used for quality control and mapping. Reads were aligned to Ensembl Genome Browser assembly ID: GRCz11. Count matrices were created with htseq-count v0.11.0^53^ and expression matrices were created with StringTie v1.3.4d^54^. Differential expression analysis was conducted using DESeq2-1.29.13^55^. Heatmaps were generated using the ComplexHeatmap v2.5.5 package for R. Raw and normalized count plots were created using ggPlot2 v3.3.2 in R. Enrichment term analysis of rescued genes was conducted using DAVID v6.8^56^ and plotted using ComplexHeatmap v2.5.5 in R. Functional enrichments nodes were categorized by GO: biological process, molecular function, and cellular component and/or KEGG or Reactome pathways using a false discovery rate (FDR) less than 0.05.

### Quantitative RT-PCR

RNA was extracted as described above. Quantitative RT-PCR (RT-qPCR) with reverse transcription was performed on a the CFX96 Touch Real-Time Detection system (BioRad) using the Luna Universal One-Step RT-qPCR kit (New England BioLabs) and primer sets validated in our lab (Supplemental Table 5). Universal 16S rRNA RT-qPCR primers were synthesized according to Clifford et al. 2012^57^

### Transgenic zebrafish

GFAP:GFP zebrafish Tg(*gfap*:GFP)^*mi2001*^ (Bernardos and Raymond, 2006) were kindly provided by Dr. Vincent Tropepe (U of Toronto) and treated as described above.

### Immunohistochemistry

Larvae were fixed at 2 dpf in 4% paraformaldehyde for 2 hours and then rinsed in PBS. larvae were then exposed to proteinase K (10ug ml^-1^ in PBT) for 20 minutes and rinsed again in PBS with 1% bovine serum albumin, 1% DMSO, and 0.1% TritonX-100 (PBDT). Larvae were blocked in 10% sheep serum in PBDT for 1 hour at room temperature and then incubated in mouse anti-alpha acetylated tubulin (Sigma-Aldrich Canada Ltd, Cat: T7451, Clone: 6-11B-1, 1:500) at 4°C for 48 hours. After 48 hours, larvae were rinsed 3 times in PBT and then incubated in the secondary antibody (1:1000) in blocking solution (2% sheep serum in PBDT) for 5 hours at room temperature. Following incubation, larvae were rinsed again 3 times in PBT and exposed to Hoechst counterstain (1:10,000) for 10 minutes at room temperature before being rinsed in PBS. 5-7 Larvae were mounted in 0.8% low melting point agar on glass bottomed imaging dish.

### Lateral line screening

Whole, 3dpf and 4dpf, Tubïgen larva from each treatment group were incubated in 4ug/ml Diasp (2-Di-4-Asp, Sigma) and 0.3 ug/ml DioC6 (3,3-dihexyloxacarbocyanine iodide, Sigma) in embryo medium for 5 minutes as per Valdivia et al^11^. After 5 minutes, larvae were rinsed 3 times in embryo medium, anesthetized in 0.4% tricaine and mounted in 0.8% low melting point agar containing 0.4% tricaine on glass bottomed imaging dish and immediately imaged by confocal microscopy, as above.

## Key Resource Table

| Reagent or resource type | Designation | Source | Identifier |
| --- | --- | --- | --- |
| Strain | Tg(*gfap*:GFP)<sup>mi2001</sup> |  | ZFIN ID: ZDB-ALT-060623-4 |
| Antibody | Monoclonal Anti-Acetylated Tubulin antibody produced in mouse | Sigma-Aldrich Canada Ltd | Cat: T7451, Clone: 6-11B-1 |
| Antibody | Donkey anti-mouse IgG (H+L) Alexa Fluor 594 | Thermo Fisher Scientific | A-2120; RRID AB_141633 |
| Vital dye | 2-Di-4-Asp | Sigma-Aldrich Canada Ltd | Cat: D3418 |
| Vital dye | 3,3-dihexyloxacarbocyanine iodide | Sigma-Aldrich Canada Ltd | Cat: 318426 |

## Supplemental tables and figures cited in text

1. Supplemental Table 1: List of 354 DE genes (downregulated in GF-CV)

2. Supplemental Table 2: GO output

3. Supplemental Table 3: David output

4. Supplemental Table 4: List of DE genes that are also SFARI genes

5. Supplemental Table 5: Excel doc of primer set sequences and efficiencies

6. Supplemental Figure 1: WMISH of 4 and 5 dpf embryos with *notch, ascl1a* and *isl1*

7. Supplemental Figure 2: 3dpf lateral line composite trunk

8. Supplemental Figure 3: 3dpf lateral line composite tail

9. Supplemental Figure 4: 4dpf lateral line composite

10. Supplemental Figure 5: Single layer composite of axonal tracks

11. Supplemental Figure 6: Projected images of a-tubulin and GFAP-GFP expression

12. Supplemental Figure 7: KEGG images

13. Supplemental Figure 8: Tail touch assay

## References

1. McFall-Ngai, M. et al. Animals in a bacterial world, a new imperative for the life sciences. Proceedings of the National Academy of Sciences 110, 3229–3236 (2013).

2. Rowland, I. et al. Gut microbiota functions: metabolism of nutrients and other food components. European Journal of Nutrition vol. 57 1 (2018).

3. Vuong, H. E. et al. The maternal microbiome modulates fetal neurodevelopment in mice. Nature 586, 281–286 (2020).

4. Erny, D. et al. Host microbiota constantly control maturation and function of microglia in the CNS. Nature Neuroscience 18, 965–977 (2015).

5. Sgritta, M. et al. Mechanisms Underlying Microbial-Mediated Changes in Social Behavior in Mouse Models of Autism Spectrum Disorder. Neuron 101, 246-259.e6 (2019).

6. Stilling, R. M. et al. Social interaction-induced activation of RNA splicing in the amygdala of microbiome-deficient mice. eLife 7, 1–21 (2018).

7. Zhu, S. et al. The progress of gut microbiome research related to brain disorders. Journal of Neuroinflammation vol. 17 25 (2020).

8. Martin, C. R., Osadchiy, V., Kalani, A. & Mayer, E. A. The Brain-Gut-Microbiome Axis. CMGH vol. 6 133–148 (2018).

9. Carabotti, M., Scirocco, A., Maselli Maria, A. & Severi, C. The gut-brain axis: interactions between enteric microbiota, central and enteric nervous systems. Annals of Gastroenterology 28, 203–209 (2015).

10. Silva, Y. P., Bernardi, A. & Frozza, R. L. The Role of Short-Chain Fatty Acids From Gut Microbiota in Gut-Brain Communication. Frontiers in Endocrinology vol. 11 25 (2020).

11. Valdivia, L. E. et al. Lef1-dependent Wnt/β-catenin signalling drives the proliferative engine that maintains tissue homeostasis during lateral line development. Development 138, 3931–3941 (2011).

12. Pujol-Martí, J. & López-Schier, H. Developmental and architectural principles of the lateral-line neural map. Frontiers in Neural Circuits 7, 47 (2013).

13. Mulligan, K. A. & Cheyette, B. N. R. Wnt signaling in vertebrate neural development and function. Journal of Neuroimmune Pharmacology vol. 7 774–787 (2012).

14. Rogan, M. R., Patterson, L. L., Wang, J. Y. & McBride, J. W. Bacterial manipulation of wnt signaling: A host-pathogen tug-of-wnt. Frontiers in Immunology vol. 10 2390 (2019).

15. Sun, J. et al. Crosstalk between NF-κB and β-catenin pathways in bacterial-colonized intestinal epithelial cells. American Journal of Physiology-Gastrointestinal and Liver Physiology 289, G129–G137 (2005).

16. Duan, Y. et al. β-Catenin activity negatively regulates bacteria-induced inflammation. Laboratory Investigation 87, 613–624 (2007).

17. Sun, J., Hobert, M. E., Rao, A. S., Neish, A. S. & Madara, J. L. Bacterial activation of β-catenin signaling in human epithelia. American Journal of Physiology-Gastrointestinal and Liver Physiology 287, G220–G227 (2004).

18. Vikström, E., Bui, L., Konradsson, P. & Magnusson, K. E. The junctional integrity of epithelial cells is modulated by Pseudomonas aeruginosa quorum sensing molecule through phosphorylation-dependent mechanisms. Experimental Cell Research 315, 313–326 (2009).

19. Kintner, J., Moore, C. G., Whittimore, J. D., Butler, M. & Hall, J. v. Inhibition of Wnt Signaling Pathways Impairs Chlamydia trachomatis Infection in Endometrial Epithelial Cells. Frontiers in Cellular and Infection Microbiology 7, 501 (2017).

20. Kessler, M. et al. Chlamydia trachomatis disturbs epithelial tissue homeostasis in fallopian tubes via paracrine Wnt signaling. American Journal of Pathology 180, 186–198 (2012).

21. Flores, R. & Zhong, G. The Chlamydia pneumoniae inclusion membrane protein Cpn1027 interacts with host cell Wnt signaling pathway regulator cytoplasmic activation/ proliferation-associated protein 2 (Caprin2). PLoS ONE 10, e0127909 (2015).

22. Cott, C. et al. Pseudomonas aeruginosa lectin LecB inhibits tissue repair processes by triggering β-catenin degradation. Biochimica et Biophysica Acta - Molecular Cell Research 1863, 1106–1118 (2016).

23. Cheesman, S. E., Neal, J. T., Mittge, E., Seredick, B. M. & Guillemin, K. Epithelial cell proliferation in the developing zebrafish intestine is regulated by the Wnt pathway and microbial signaling via Myd88. Proceedings of the National Academy of Sciences of the United States of America 108, 4570–4577 (2011).

24. Neumann, P. A. et al. Gut commensal bacteria and regional Wnt gene expression in the proximal versus distal colon. American Journal of Pathology 184, 592–599 (2014).

25. Chen, B. et al. Small molecule-mediated disruption of Wnt-dependent signaling in tissue regeneration and cancer. Nature Chemical Biology 5, 100–107 (2009).

26. Huang, S. M. A. et al. Tankyrase inhibition stabilizes axin and antagonizes Wnt signalling. Nature 461, 614–620 (2009).

27. Meijer, L. et al. GSK-3-Selective Inhibitors Derived from Tyrian Purple Indirubins. Chemistry and Biology 10, 1255–1266 (2003).

28. Xie, Y. et al. Lef1-dependent hypothalamic neurogenesis inhibits anxiety. PLoS Biology 15, e2002257 (2017).

29. Schredelseker, T. & Driever, W. Conserved Genoarchitecture of the Basal Hypothalamus in Zebrafish Embryos. Frontiers in Neuroanatomy 14, 3 (2020).

30. Zarrinpar, A. et al. Antibiotic-induced microbiome depletion alters metabolic homeostasis by affecting gut signaling and colonic metabolism. Nature Communications 9, 1–13 (2018).

31. Sewell, D. L., Wostmann, B. S., Gairola, C. & Aleem, M. I. H. Oxidative energy metabolism in germ free and conventional rat liver mitochondria. American Journal of Physiology 228, 526–529 (1975).

32. Luczynski, P. et al. Growing up in a bubble: Using germ-free animals to assess the influence of the gut microbiota on brain and behavior. International Journal of Neuropsychopharmacology 19, 1–17 (2016).

33. Bäckhed, F. et al. The gut microbiota as an environmental factor that regulates fat storage. Proceedings of the National Academy of Sciences of the United States of America 101, 15718–23 (2004).

34. Ganz, J., Melancon, E. & Eisen, J. S. Zebrafish as a model for understanding enteric nervous system interactions in the developing intestinal tract. Methods in Cell Biology 134, 139–164 (2016).

35. Schretter, C. E. et al. A gut microbial factor modulates locomotor behaviour in Drosophila. Nature vol. 563 402–406 (2018).

36. Lush, M. E. & Piotrowski, T. Sensory hair cell regeneration in the zebrafish lateral line. Developmental Dynamics vol. 243 1187–1202 (2014).

37. Clingman, C. C. & Ryder, S. P. Metabolite sensing in eukaryotic mRNA biology. Wiley Interdisciplinary Reviews: RNA 4, 387–396 (2013).

38. Bhat, M. I. & Kapila, R. Dietary metabolites derived from gut microbiota: Critical modulators of epigenetic changes in mammals. Nutrition Reviews 75, 374–389 (2017).

39. Nankova, B. B., Agarwal, R., MacFabe, D. F. & la Gamma, E. F. Enteric Bacterial Metabolites Propionic and Butyric Acid Modulate Gene Expression, Including CREB-Dependent Catecholaminergic Neurotransmission, in PC12 Cells - Possible Relevance to Autism Spectrum Disorders. PLoS ONE 9, e103740 (2014).

40. Navajas Acedo, J. et al. PCP and Wnt pathway components act in parallel during zebrafish mechanosensory hair cell orientation. Nature Communications 10, 1–17 (2019).

41. Bem, J. et al. Wnt/β-catenin signaling in brain development and mental disorders: keeping TCF7L2 in mind. FEBS Letters vol. 593 1654–1674 (2019).

42. Amoyel, M., Cheng, Y. C., Jiang, Y. J. & Wilkinson, D. G. Wnt1 regulates neurogenesis and mediates lateral inhibition of boundary cell specification in the zebrafish hindbrain. Development 132, 775–785 (2005).

43. Phelps, D. et al. Microbial colonization is required for normal neurobehavioral development in zebrafish. Scientific Reports 7, (2017).

44. Davis, D. J., Bryda, E. C., Gillespie, C. H. & Ericsson, A. C. Microbial modulation of behavior and stress responses in zebrafish larvae. Behavioural Brain Research 311, 219–227 (2016).

45. Ghanavati, R. et al. Lactobacillus species inhibitory effect on colorectal cancer progression through modulating the Wnt/β-catenin signaling pathway. Molecular and Cellular Biochemistry 470, 1–13 (2020).

46. Silva-García, O., Valdez-Alarcón, J. J. & Baizabal-Aguirre, V. M. Wnt/β-catenin signaling as a molecular target by pathogenic bacteria. Frontiers in Immunology vol. 10 (2019).

47. Liu, X. et al. Wnt2 inhibits enteric bacterial-induced inflammation in intestinal epithelial cells. Inflammatory Bowel Diseases 18, 418–429 (2012).

48. Westerfield, M. The Zebrafish Book. A Guide for the Laboratory Use of Zebrafish (Danio rerio). (Eugene, OR, Uinversity of Oregon Press, 1995).

49. Thisse, C. & Thisse, B. High-resolution in situ hybridization to whole-mount zebrafish embryos. Nature Protocols 3, 59–69 (2008).

50. Collymore, C. Anesthesia, analgesia, and euthanasia of the laboratory zebrafish. in The Zebrafish in Biomedical Research: Biology, Husbandry, Diseases, and Research Applications 403–413 (Elsevier, 2019). doi:10.1016/B978-0-12-812431-4.00034-8.

51. Pertea, M., Kim, D., Pertea, G. M., Leek, J. T. & Salzberg, S. L. Transcript-level expression analysis of RNA-seq experiments with HISAT, StringTie and Ballgown. Nature Protocols 11, 1650–1667 (2016).

52. Kim, D., Langmead, B. & Salzberg, S. L. HISAT: A fast spliced aligner with low memory requirements. Nature Methods 12, 357–360 (2015).

53. Anders, S., Pyl, P. T. & Huber, W. HTSeq - A Python framework to work with high-throughput sequencing data. bioRxiv 002824 (2014) doi:10.1101/002824.

54. Pertea, M. et al. StringTie enables improved reconstruction of a transcriptome from RNA-seq reads. Nature Biotechnology 33, 290–295 (2015).

55. Love, M. I., Huber, W. & Anders, S. Moderated estimation of fold change and dispersion for RNA-seq data with DESeq2. Genome Biology 15, 550 (2014).

56. Huang, D. W., Sherman, B. T. & Lempicki, R. A. Systematic and integrative analysis of large gene lists using DAVID bioinformatics resources. Nature Protocols 4, 44–57 (2009).

57. Clifford, R. J. et al. Detection of bacterial 16S rRNA and identification of four clinically important bacteria by real-time PCR. PloS one 7, e48558 (2012).

