## Supplemental Figures for "Gut-derived metabolites influence neurodevelopmental gene expression and Wnt signalling events in a germ-free zebrafish model"

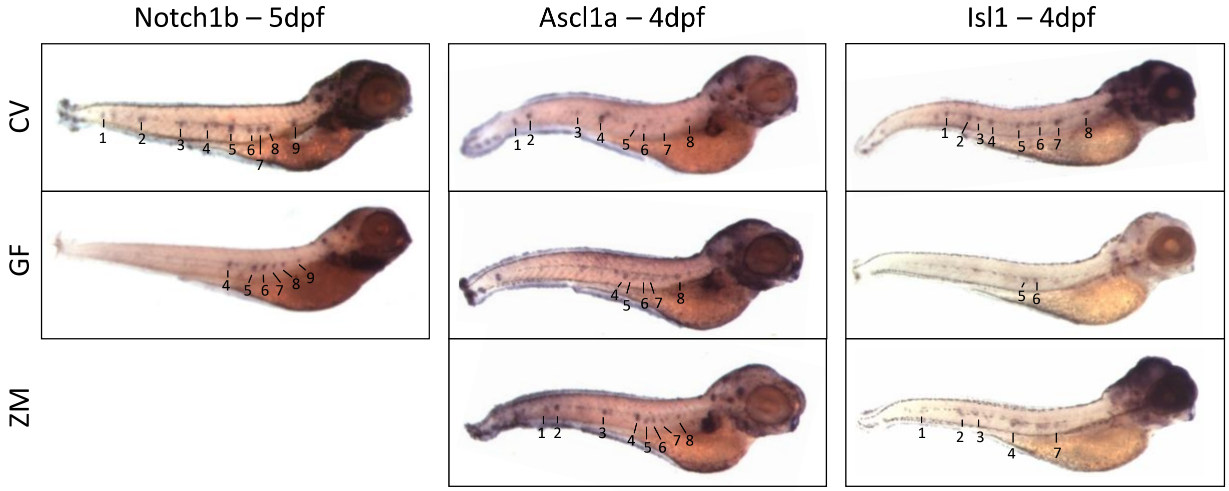


**Supplemental Figure 1.** Representative images of whole mount *in situ* hybridization on 5 dpf conventional embryos and germ-free embryos with *Notch1b* and 4 dpf CV, GF and ZM embryos with *acsl1a* and *Isl1*.

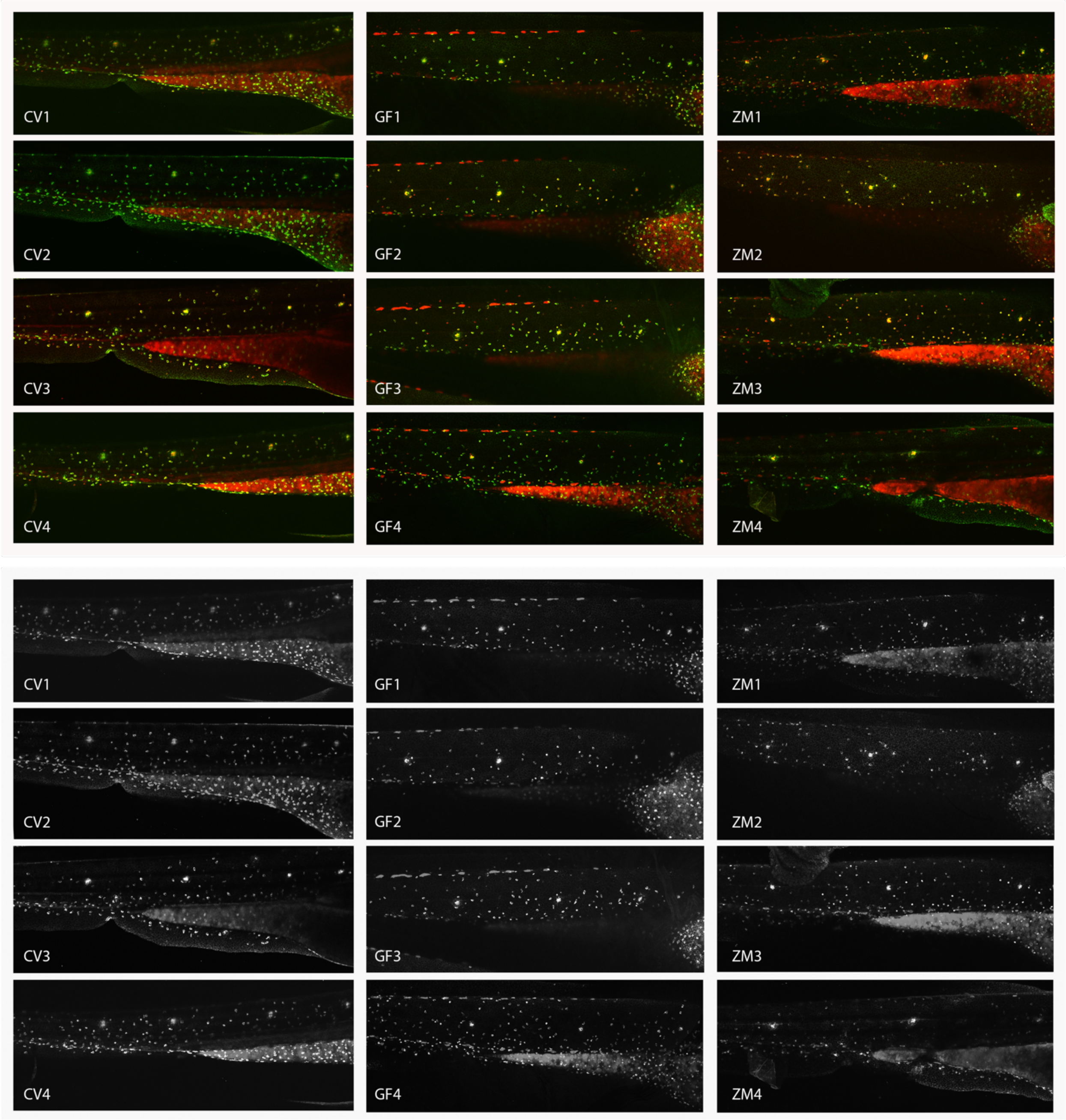


**Supplemental figure 2.** Composite images of 3dpf embryos incubated in a mixture of Diasp and DioC6 to mark neuromast hair and accessory cells of the posterior lateral line. Bottom image is composed of same images as the top but in greyscale.

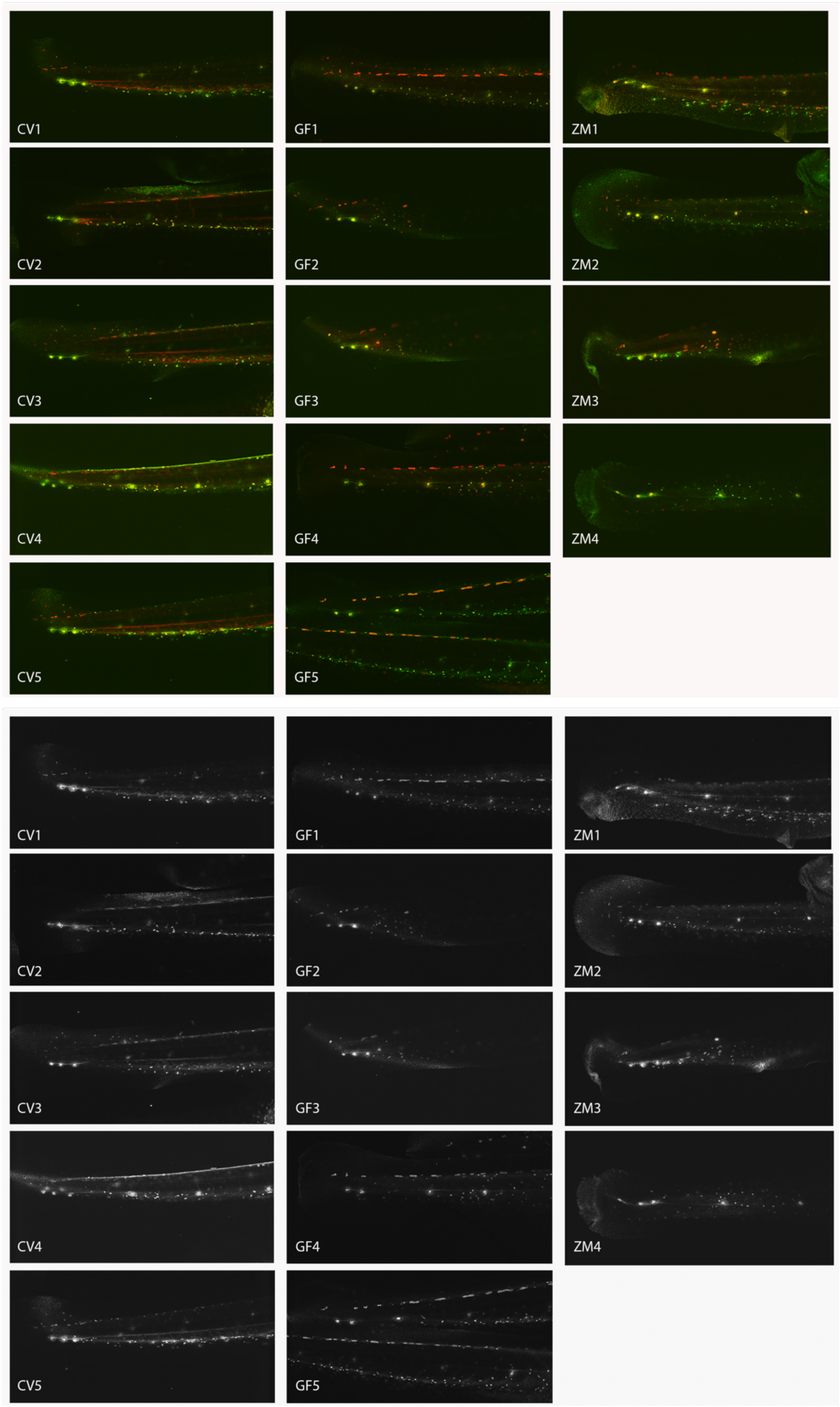


**Supplemental figure 3.** Composite images of 3dpf embryos incubated in a mixture of Diasp and DioC6 to mark neuromast hair and accessory cells of the posterior lateral line. Images show the tail bud, with terminal neuromasts and posterior neuromasts of the trunk. Bottom image is composed of same images as the top but in greyscale.

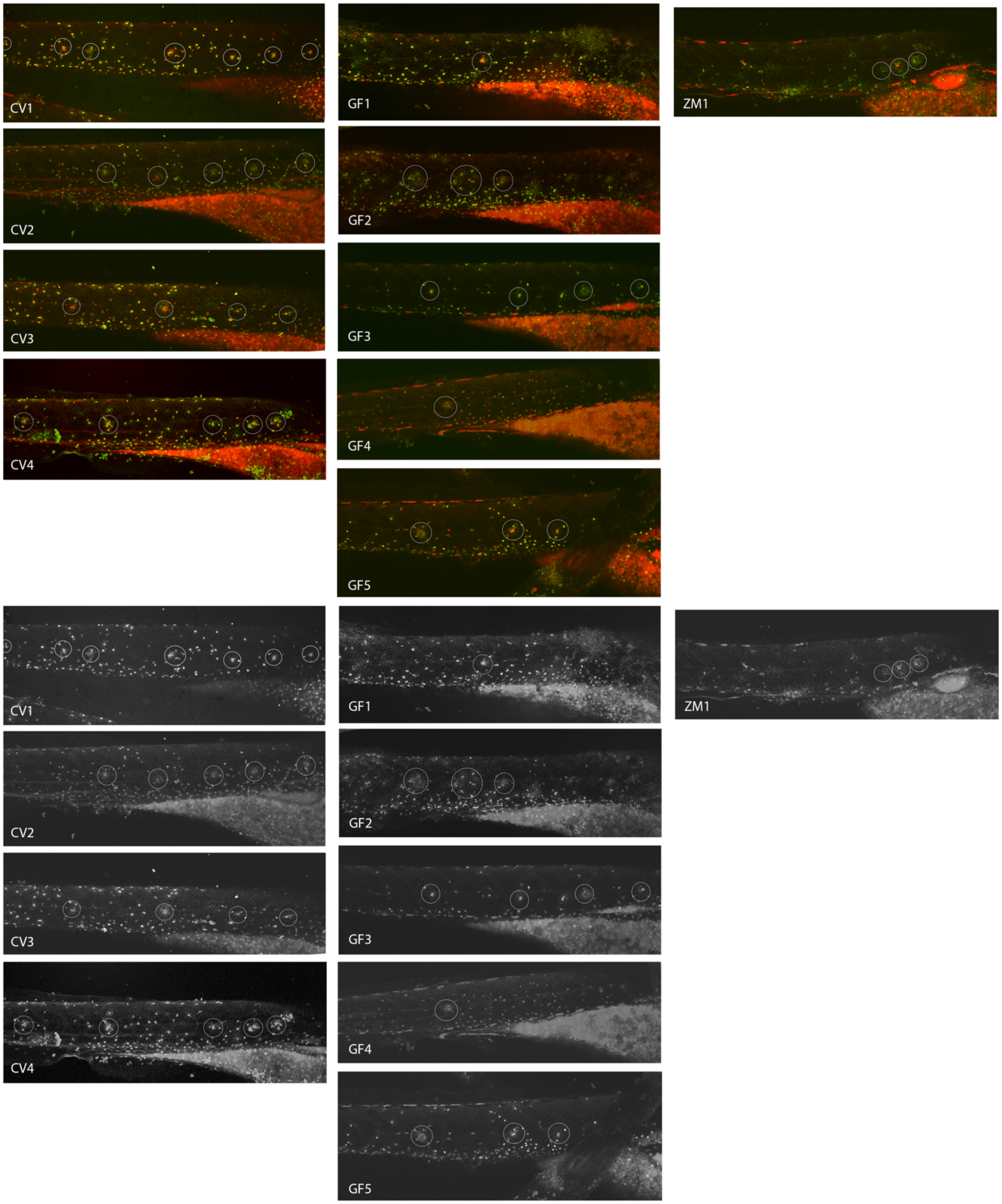


**Supplemental figure 4**. Composite images of 4dpf embryos in a mixture of Diasp and DioC6 to mark neuromast hair and accessory cells of the posterior lateral line. The CV embryos have an average of 5 neuromasts (range from 4-7), whereas GF embryos have less on average (range from 1-4). Bottom image is composed of same images as the top but in greyscale.

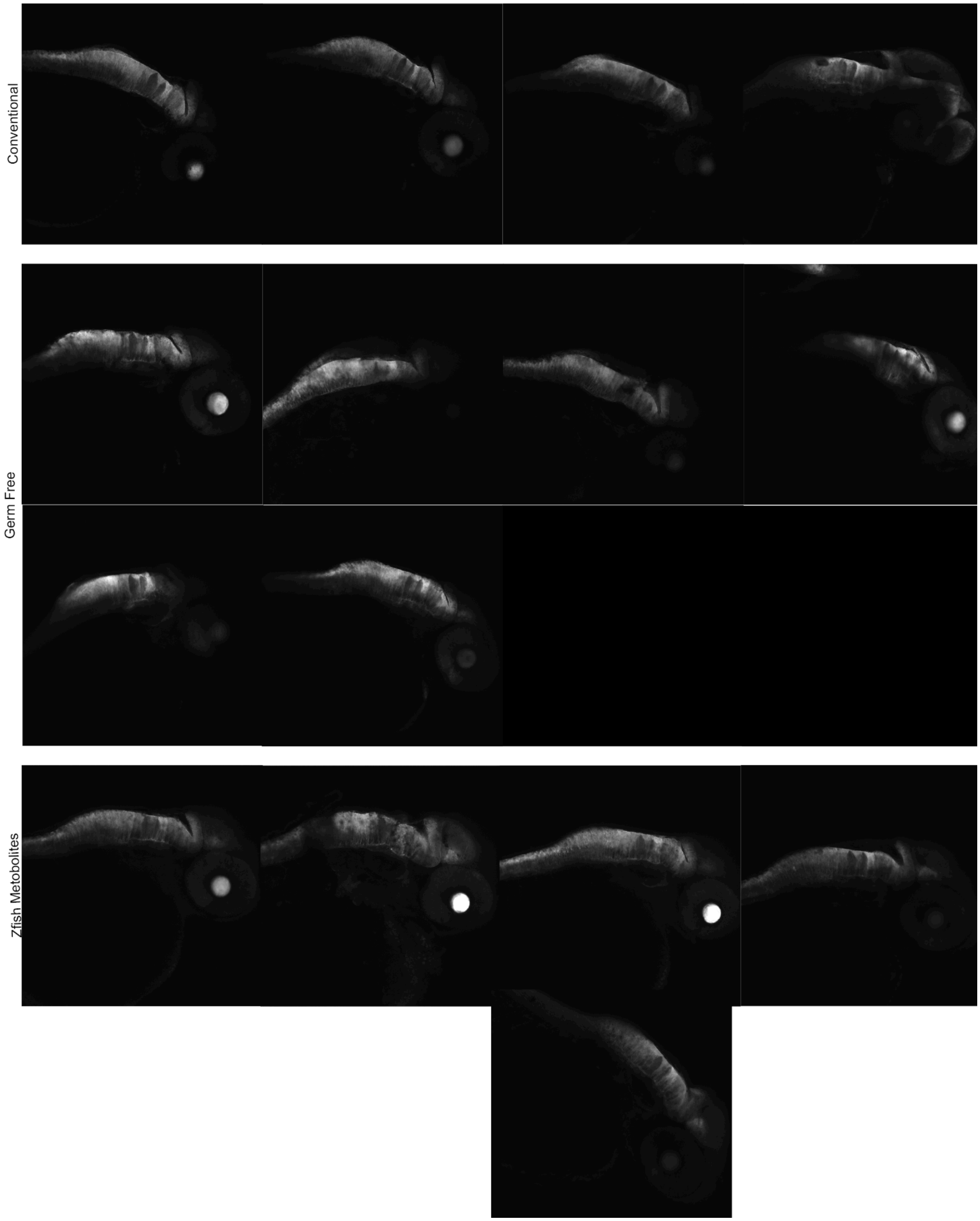


**Supplementary Figure 5.** Single layer composite using the Axonal tracts in the hindbrain as a guide.

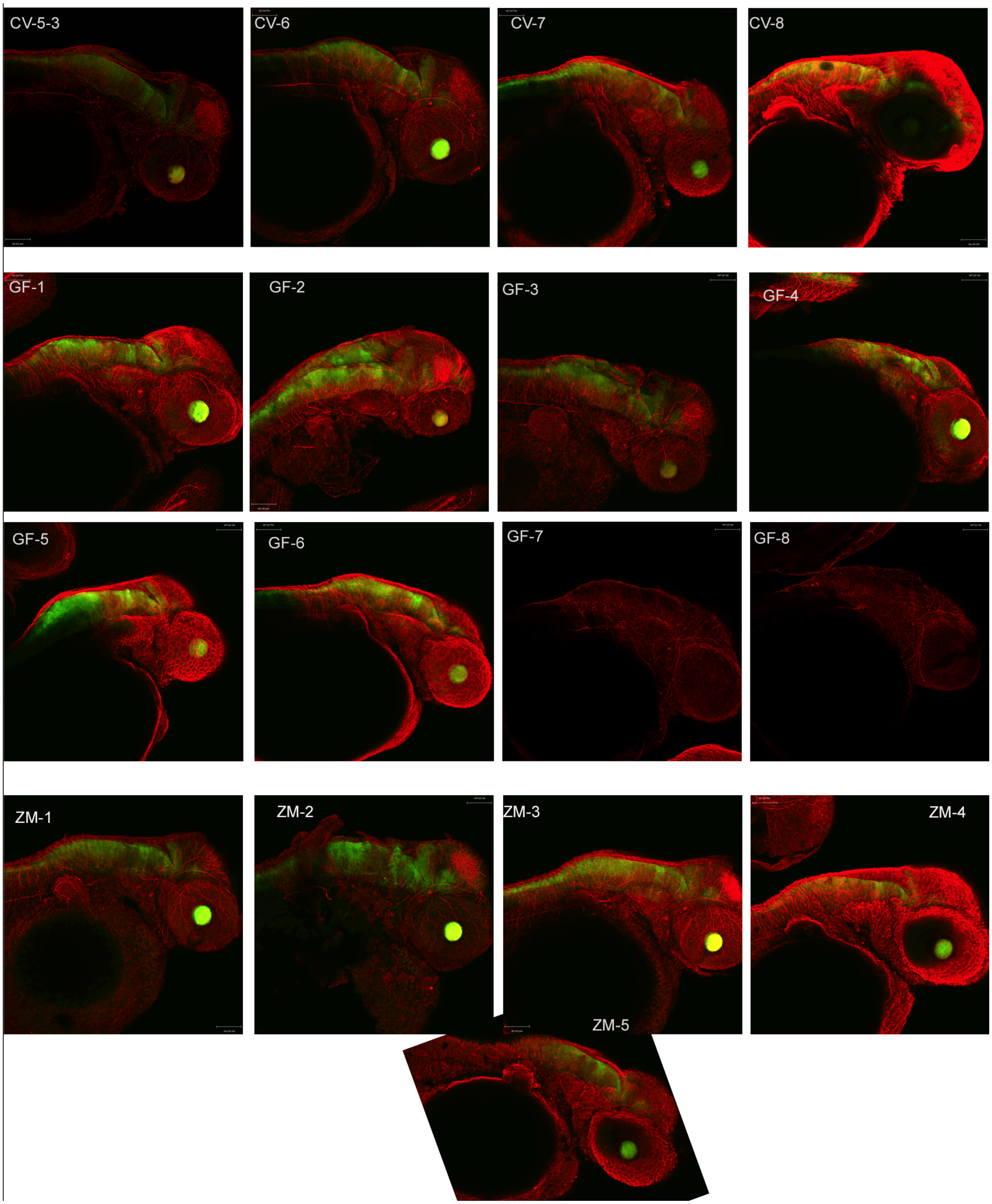


**Supplementary Figure 6**. Projected images of a-tubulin and GFAP-GFP expression.

### Signalling pathways:

A

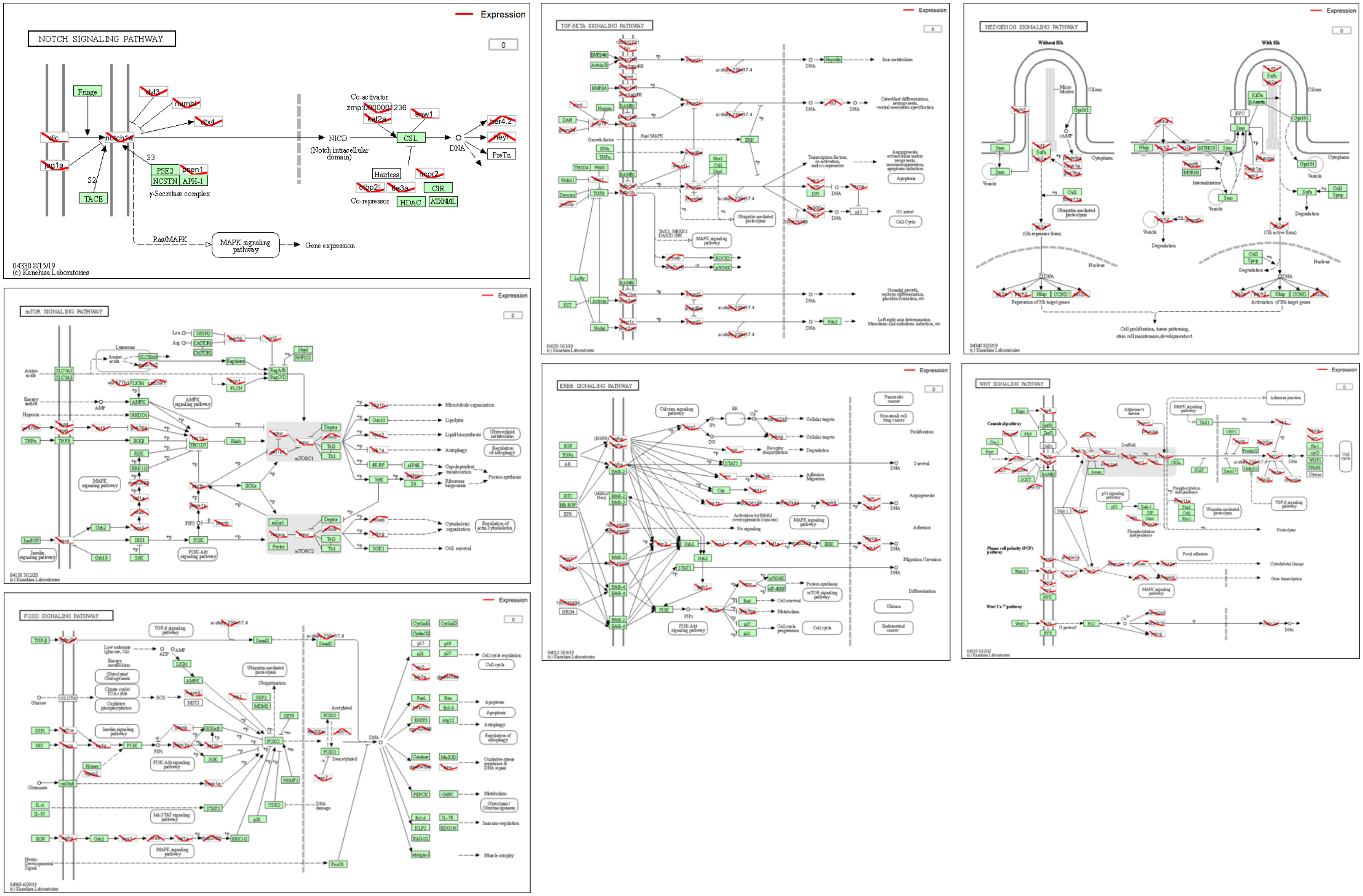

RNA associated:

B

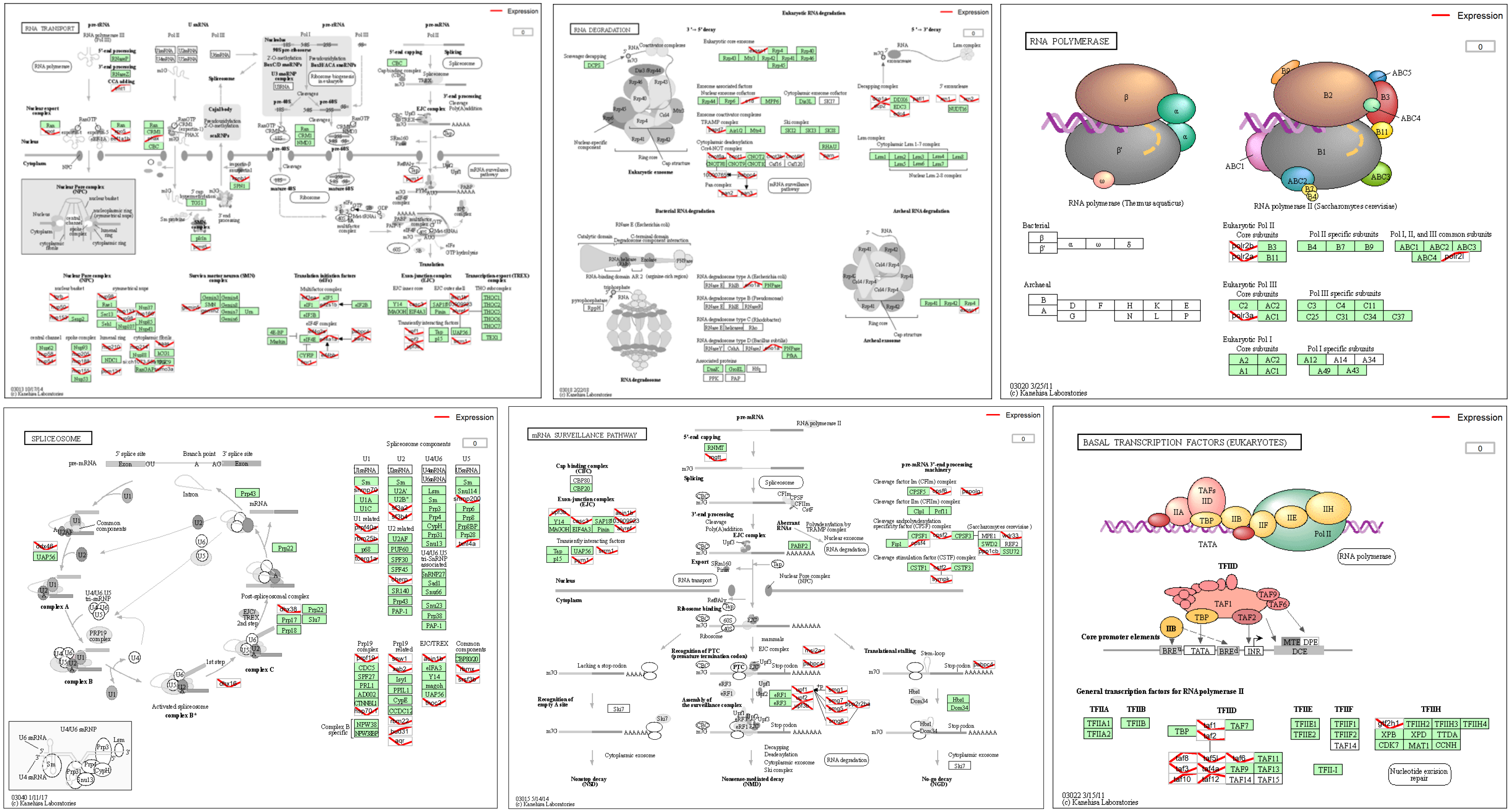

Other:

C

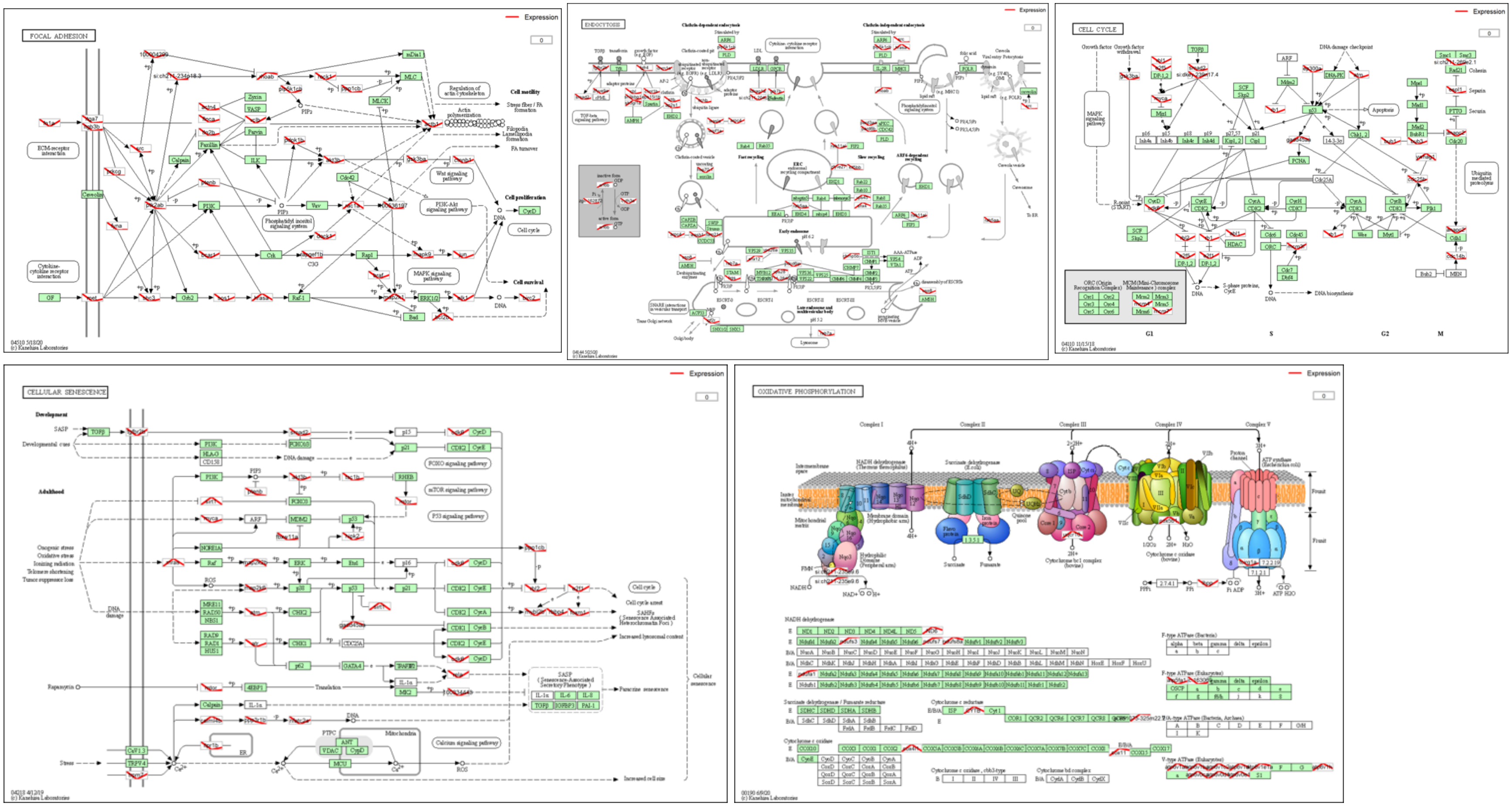


**Supplemental Figure 7.** KEGG profile output of important pathways and genes from complete RNA-Seq dataset. A) Signalling pathways, B) RNA associated pathways and ontology, and C) other. Levels are shown in red where the left side (CV) is arbitrarily set to 0, the middle point is GF and the right point is ZM.

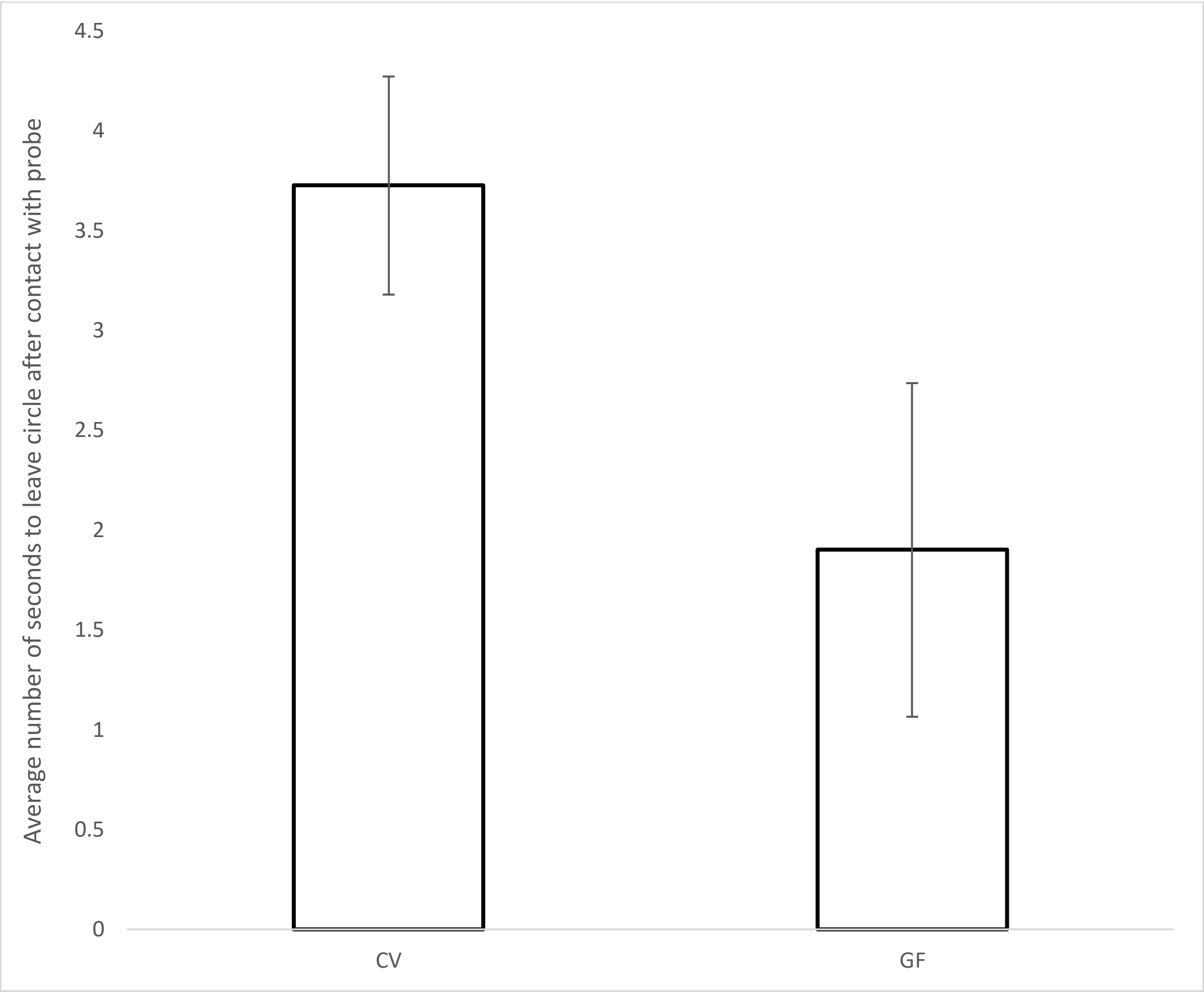


**Supplemental Figure 8. Preliminary tail touch assay.** Bar graph displaying the average number of seconds for 3dpf CV (N=7) and GF (N=9) embryos to swim out of a designated area after contact with a probe. P = 0.078228 in a standard weighted means analysis one-way ANOVA with 2 independent variables. Videos were recorded at 500 frames per second and calculations were made using ImageJ.
